# Theranostic Bottle-Brush Polymers Tailored for Universal Solid-Tumour Targeting

**DOI:** 10.1101/2023.07.13.548666

**Authors:** Wei Zhang, Yanwen Xu, Rongjun Guo, Peiling Zhuang, Huixia Hong, Hui Tan, Mingfeng Wang

## Abstract

Nanomedicines involving nanotechnologies and engineering of nanomaterials for medicines have shown great promise in diagnosis and treatment of diseases including cancers. A major hurdle that limits the successful clinical translation of nanomedicines, however, is how to overcome the cascaded biological barriers and improve the delivery efficacy towards the disease sites and minimize the toxicity against healthy tissues and cells. Here, we report a type of bottle-brush-like polymers systematically optimized in their chemical structures, sizes, and surface charges that lead to their outstanding pharmacokinetics and tumour-targeting performances in a variety of both subcutaneous and orthotopic tumour models. The potential mechanism has been studied by revealing the structure-activity relationship of these polymers in overcoming the biological barriers, including their avoidance by the immune system and deep tumour infiltration. Our study may offer insight for a rational design of highly efficient delivery platform of polymeric nanomedicines that could effectively overcome the cascaded biological barriers and thus lead to high tumour-targeting efficacy and low toxicity.

## Introduction

Nanomedicines have played an important role in both diagnosis and therapy of a variety of cancers^1-9^. However, successful translation of cancer nanomedicines remains rare despite decades of intensive research^10-12^. A major hurdle towards clinical translation is that most nanomedicines have to experience a series of biological barriers before reaching tumour sites^13-17^. Generally, intravascularly administrated nanoparticles (NPs) are immediately adsorbed by plasma proteins (namely opsonization), significantly altering their surface properties and in vivo trajectory. Then majority of the NPs with a size larger than 10 nm are cleared by liver and spleen of the reticuloendothelial system (RES), whereas NPs small than 6 nm are often filtered out by kidney. The remaining NPs must extravasate through the tumour vasculature and penetrate through the dense extracellular matrix to reach the targeted cells and release the cargos/drugs (Fig. 1c)^18,19^. As a consequence, a statistical survey revealed that only 0.7% (median) of the injected dose of NPs is delivered to solid tumour^2,20^. Up to 99% of the NPs are eliminated in the abovementioned biological barriers, particularly by mononuclear phagocytic system (MPS) and renal system^20^. Furthermore, the enhanced permeability and retention (EPR) effect varies significantly with locations, stages and sizes of tumours, patients and cancer types due to the heterogenous pathophysiological properties (vascular density and leakiness, interstitial fluid pressure, etc.) of tumour microenvironment^21^, making it complicated to design a universal and efficient nanocarrier. Thus, rational design of nanocarriers is urgently needed to overcome these biological barriers and improve the efficacy of targeting delivery to a wide variety of tumours.

**Figure 1.**
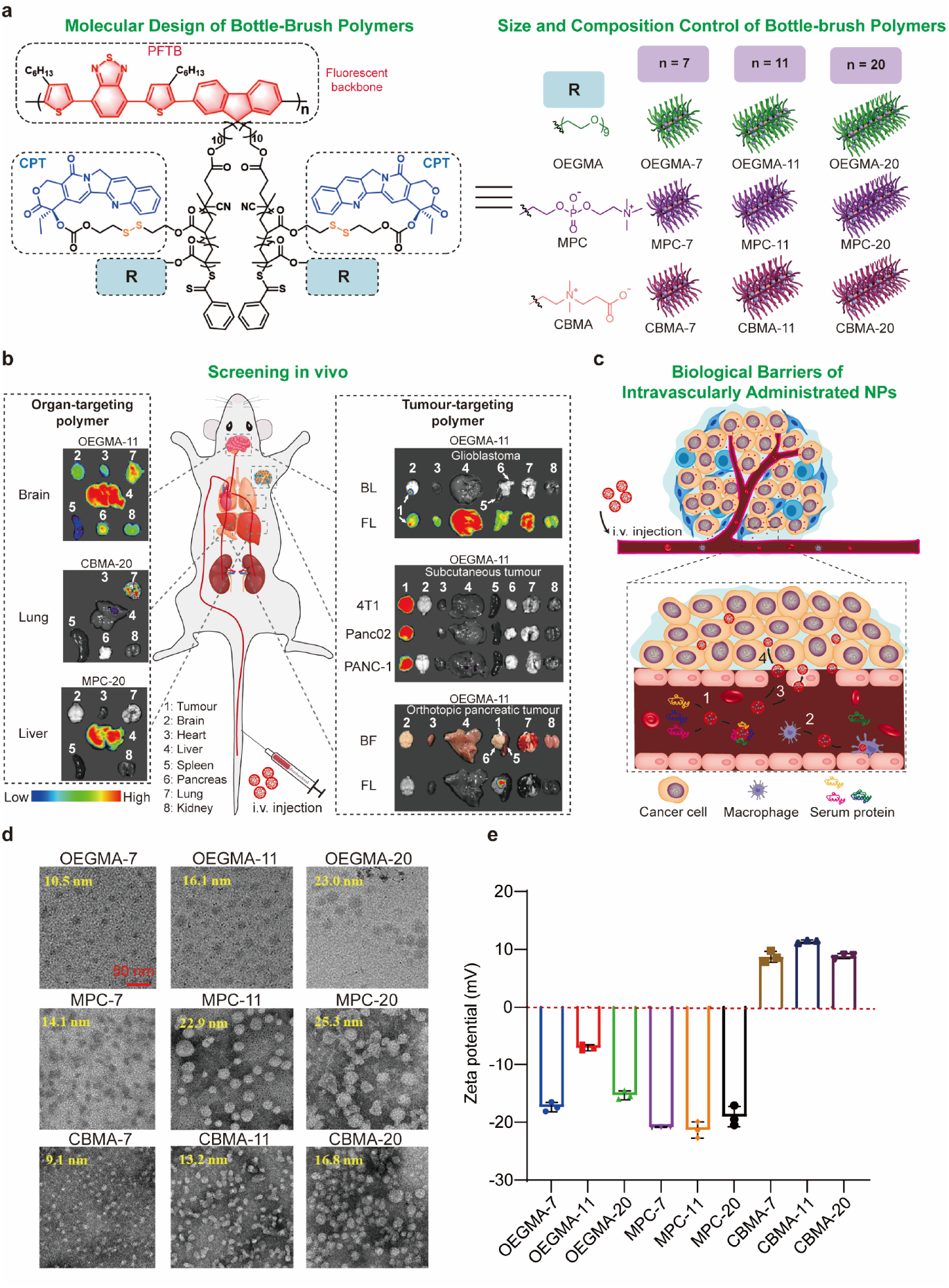
Molecular design and characterization of theranostic bottle-brush polymers and the as-formed NPs for specific tumour- and tissue-targeting delivery. **a** The chemical structures and schematic view of polymers. Pink column denotes the fluorescent backbone of bottle-brush polymer; Blue sphere denotes CPT; green, purple and orange strands denote the side chains polymerized by OEGMA, MPC and CBMA. For instance, the bottle-brush polymer named OEGMA-7 is composed of the backbone with 7 repeating units and the side chain polymerized by OEGMA monomer and CPT prodrug. **b** Screening in several tumour models and biodistribution profiles of polymeric NPs: OEGMA-11 NPs was able to pass through blood brain barrier (BBB) and effectively accumulate in different tumour models; CBMA-20 and MPC-20 NPs can target lung and liver, respectively. BL, bioluminescence; FL, fluorescence; BF, bright field; i.v., intravenous. **c** Schematic illustration of journey that intravenously administered NPs experienced: 1. the formation of protein corona on the surface of NPs, 2. the internalization of NPs by macrophages, 3. NPs extravasating through tumour vasculature, 4. NPs penetrating in tumour tissues. **d, e** Transmission electron microscopy (TEM) images (**d**) and zeta potential (**e**) of NPs in water (n = 3). The average diameters of the polymeric NPs labeled on the TEM images (**d**) were measured from at least 50 NPs using Image J. Data were presented as mean ± s.d..

The physicochemical properties of NPs (size, charge, shape, rigidity, surface chemistry, etc.) determine their fates in vivo^2,19,22,23^. Polymers with excellent control over their structural and functional compositions provide a promising platform to develop customized drug-delivery vehicles. For instance, to cloak NPs and prolong their blood retention time, PEGylation has been widely utilized as a gold standard in the design of drug carriers^24^. Coating NPs with zwitterion groups, such as 2-methacryloyloxyethyl phosphorylcholine (MPC) and carboxybetaine methacrylate (CBMA)^25,26^, also minimized opsonization and improved pharmacokinetics. In addition, extensive studies verified the relevance between size and in vivo performance of NPs^27,28^.

Despite some advances on the development of new materials and technologies to enhance the tumour-targeting efficacy of surface-modified polymers^4-7,12,27,29-33^, and mesoporous silica NPs (Table S1, Supporting Information)^34,35^, there remains no rational design principle yet for nanomedicines that integrate factors of both structures and physicochemical properties (such as sizes, shapes, surface charges, hydrophobic/hydrophilic ratios, and rigidities), in order to overcome the cascaded biological barriers that most intravenously administered nanomedicines have to face before reaching the tumour sites in an efficient way. Here, we report such a type of bottle-brush-like polymers systematically optimized in their chemical structures, sizes, rigidities, and surface charges that lead to their outstanding pharmacokinetics and tumour-targeting performances in a variety of both subcutaneous and orthotopic tumour models.

Fig. 1a depicts the molecular design of our bottle-brush polymers. All the polymers bare the same type of relatively rigid backbone with π-conjugated structures and strong red-fluorescent properties that benefit the imaging and tracking of the polymers both in vivo and in vitro. The chain lengths of the fluorescent backbones are tunable by changing the number-average degree (n) of polymerization from 7 to 11 to 20. A series of polymer chains containing hydrophilic monomers, including (oligo(ethylene glycol) methyl ether methacrylate) (OEGMA), CBMA, MPC were grafted together with a hydrophobic monomer of camptothecin (CPT) as an anticancer prodrug from the fluorescent backbone via controlled free-radical polymerization. The resulting series of bottle-brush-like polymers, denoted as OEGMA-(7,11,20), MPC-(7,11,20), and CBMA-(7,11,20), respectively, have distinct sizes and surface properties that strongly relate to their biological fate in vivo.

Our systematic screening of these theranostic polymers, which integrate both imaging and therapeutic components in one polymer, in several subcutaneous and orthotopic tumour models, (Fig. 1b) has led to exceptional tumour-targeting efficiency of OEGMA-11 polymer as well as their brain-penetrating capability, whereas MPC and CBMA NPs surprisingly accumulate most in the liver and lung but poor tumour-targeting performances. To understand the mechanism that may lead to the exceptional pharmacokinetics and tumour-targeting efficacy of OEGMA-11 polymer and the as-formed NPs, we have characterized the behaviors of the NPs in overcoming the biological barriers, including their avoidance by the immune system and deep tumour infiltration (Fig. 1c). Our results may offer insight for a rational design of highly efficient delivery platform of polymeric nanomedicines that could effectively overcome the cascaded biological barriers and thus lead to high tumour-targeting efficacy and low toxicity.

## Results

### Design, synthesis and characterization of polymers

Controlling the reaction time of polycondensation between 4,7-bis(4-hexylthien)-2,1,3-benzo-thiadiazole and 2,2’-(((2,7-dibromo-9H-fluorene-9,9-diyl)bis(undecane-11,1-diyl))bis(oxy))bis(tetrahydro-2H-pyran) as monomers via a scheme of C-H direct arylation led to the fluorescent backbones ((poly(fluorene-*alt*-(4,7-bis(hexylthien)-2,1,3-benzothiadiazole)) (PFTB)) with tunable average numbers (n = 7, 11 and 20) of repeating units^29,36-38^, from which the side chains were grafted by copolymerization of camptothecin (CPT) prodrug and OEGMA/MPC/CBMA monomer at a molar ratio of 1:5. In this synthetic protocol, the chemotherapeutic components of CPT were conjugated to polymer through the glutathione-responsive disulfide bonds, which allowed CPT to be controllably released in glutathione-enriched cancer cells^39-41^.

The detailed synthetic schemes are described in Scheme S1-S2. The structures and repeating units of PFTB backbones were confirmed by proton nuclear magnetic resonance (^1^H-NMR) spectroscopy (Figure S1) and gel permeation chromatograph (GPC) (Table S2). Then the structures of OEGMA polymers were verified by ^1^H-NMR (Fig. S4), while Fourier transform infrared (FTIR) spectroscopy was conducted to characterize the structures of CBMA and MPC polymers (Fig. S5, S6) due to their poor solubilities in a single organic solvent, indicating MPC and CBMA monomers were successfully conjugated to the backbones (PFTB). In addition, the successful grafting of CPT prodrugs from the PFTB backbones was evidenced by co-presence of the absorption peaks of PFTB (517 nm) and CPT (358 nm) in the UV-Vis absorption spectra of OEGMA, MPC, CBMA NPs (Fig. S2). The variation of absorbance ratios of CPT over PFTB (Fig. S3) indicates relatively high content of CPT in polymers of OEGMA-7 and CBMA-20, while the remaining polymers appear similar to each other. All the polymers can be well dispersed into aqueous media and show strong red fluorescence under UV (365 nm) light irradiation, except OEGMA-7 and OEGMA-20 which show purple fluorescence mainly due to the relatively high content of CPT in these two polymers.

Polymeric NPs were prepared via nano-precipitation, in which an aliquot of OEGMA in tetrahydrofuran (THF), MPC in THF/ethanol or CBMA in THF/dimethylformamide, respectively, was rapidly injected into deionized water. The organic solvent was then removed by dialysis against deionized water. As shown in Fig. S2 and S7, all NPs exhibited strong fluorescence and were stable after two years of storage in water, owing to the hyperbranched structures of the polymers and the feature of unimolecular micelles^31,37,38,42-45^. The NPs show uniform spherical morphologies as observed by transmission electron microscopy (TEM, Fig. 1d). As expected, diameters of NPs gradually increased with the increase of chain length of backbone in the range of 9-25 nm, generally smaller than their hydrodynamic sizes measured by dynamic light scattering (DLS) (Fig. S7). Moreover, the surface zeta potentials were approximately - 20.5 and + 9.3 mV for MPC and CBMA NPs, respectively. Strikingly, the surface zeta potential of OEGMA-11 NPs was at - 7.0 mV, much higher than those of OEGMA-7 and -20 NPs (both around - 16.4 mV) (Fig. 1e), which could be related to the different loading amounts of CPT between OEGMA NPs.

### In vivo screening of NPs

The delivery profiles of polymeric NPs were first investigated in C57BL/6 mice bearing subcutaneous Panc02 (mouse pancreatic cancer cell line) tumour at a polymer dose of 2 mg/kg. OEGMA-(7, 11, 20), MPC-(7, 11, 20) and CBMA-(7, 11, 20) NPs were injected intravenously into mice via tail vein and mean fluorescence intensity (MFI) of main organs and tumours were measured by in vivo imaging system (IVIS) at 48 h postinjection. As shown in Fig. 2a,b and S8, OEGMA-11 NPs exhibited the highest ratio of distribution in the tumours among all organs. Relatively weak fluorescence in the tumour and the highest fluorescence in the liver were observed in the mice administrated with MPC NPs, and their accumulation in tumour decreased rapidly with increase of size. Interestingly, CBMA NPs barely accumulated in the liver and selectively distributed in the lung and spleen when the sizes of the NPs were increased.

**Figure 2.**
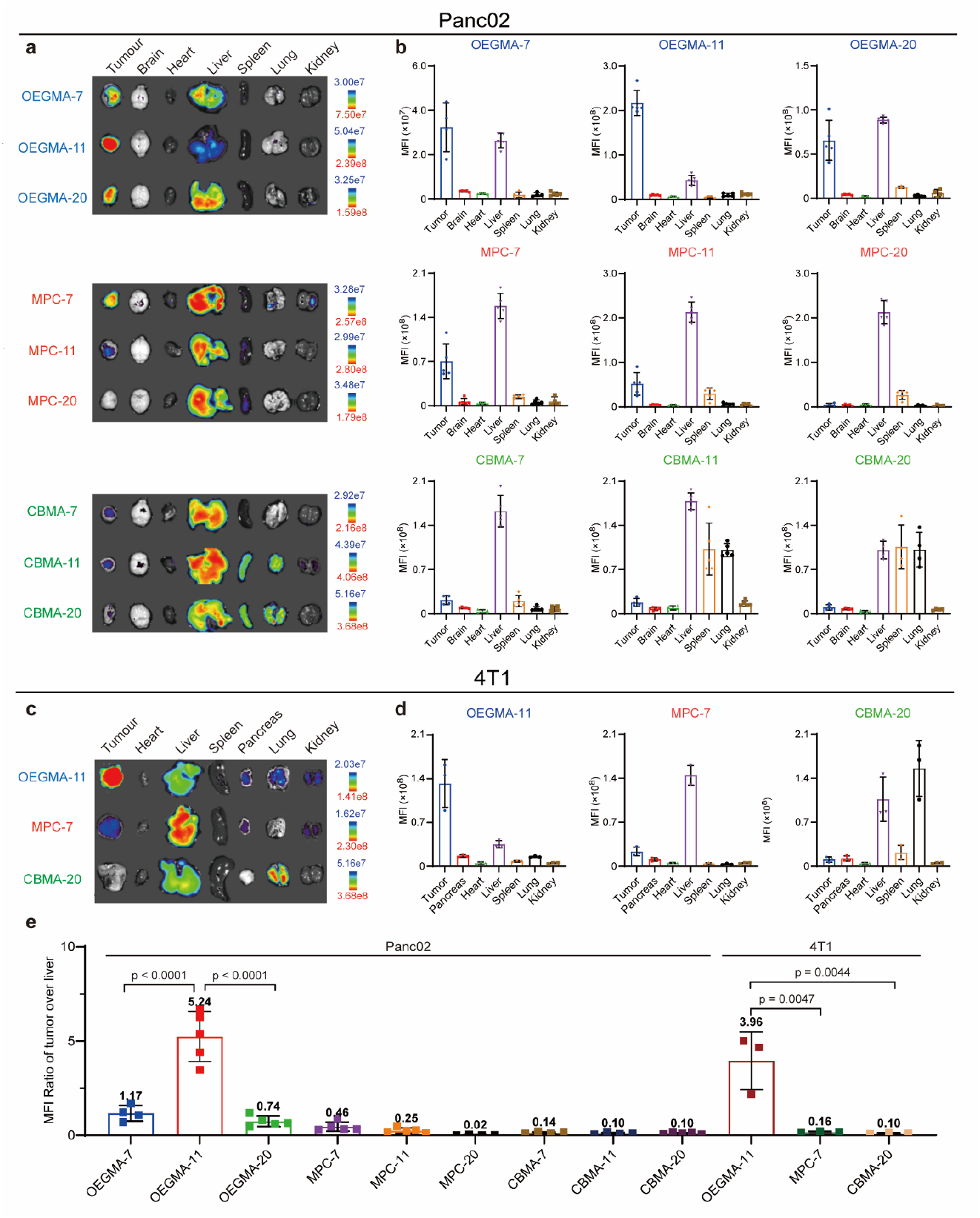
Proper size and surface properties allow polymeric NPs to selectively accumulate in organs and tumours. **a, b** Ex vivo fluorescence imaging (**a**) and mean fluorescence intensity (MFI) (**b**) of main organs and Panc02 tumours extracted from C57BL/6 at 48 h after tail-vein injection of NPs (n = 4 or 5). **c, d** Ex vivo fluorescence imaging (**c**) and MFI (**d**) of main organs and 4T1 tumours extracted from BALB/c at 48 h after tail-vein injection of NPs (n = 3). **e** MFI ratio between Panc02/4T1 tumour and the liver from **b**,**d** (n = 3 or 4 or 5). Data were presented as mean ± s.d. and statistical significance was determined using one-way ANOVA followed by Tukey’s multiple comparisons test.

To examine whether the tumour-targeting ability of our NPs was universal in other tumour models, we evaluated further the biodistribution of OEGMA-11, using MPC-7 and CBMA-20 NPs as control, in BALB/c mice bearing subcutaneous 4T1 (mouse breast cancer cell line) tumour (Fig. 2c,d & S9). Similar to Panc02 tumour model as described above, OEGMA-11 NPs showed excellent tumour-targeting again in 4T1 tumour model. The tumour-targeting efficiency of MPC-7 NPs appeared lower than that in Panc02 tumour model. CBMA-NPs showed less distribution in spleen of mice bearing 4T1 tumour compared to Panc02 tumour model. It should be noted that CBMA-20 NPs exhibited noticeable distribution in lung in both Panc01- and 4T1-bearing mice. We used the MFI ratio (denoted as R value) of the tumour over the liver to define the tumour-targeting ability of NPs.

As shown in Fig. 2e, OEGMA-11 NPs performed the highest R values (5.2 for Panc02 and 4.0 for 4T1) among all NPs, demonstrating outstanding tumour-targeting ability of OEGMA-11 NPs. Overall, OEGMA-11 NPs exhibited excellent tumour-targeting ability in subcutaneous Panc02 and 4T1 tumour models and were selected as a candidate of tumour-targeting nanocarrier for further research.

### Tumour-targeting behavior of OEGMA-11 NPs at a therapeutic dose

We further characterized behavior in vivo, particularly the time-dependent biodistribution, of OEGMA-11 NPs with a dose at therapeutic level, much higher than that for bioimaging. First, the accumulation of OEGMA-11 NPs in tumour was reevaluated in subcutaneous 4T1 and Panc02 tumour models. OEGMA-11 NPs at a dose of 40 mg/kg were injected into 4T1 tumour-bearing mice and imaging of the main organs and tumours was performed *ex vivo* on Day 1, 2, 3 postinjection. The MFI of the tumours was remarkably high compared to those organs and continued increasing over 3 days (Fig. 3a,b & S10). The average R values were 13.2 on Day 1, and exceeded 13.9 and 14.3 on Day 2 and 3, respectively, due to the strong fluorescence from the tumours beyond the measurement range of IVIS (Fig. 3c), and decreased to 7.03 on Day 2 in the bigger 4T1 tumours with the internal necrosis (Fig. S11).

**Figure 3.**
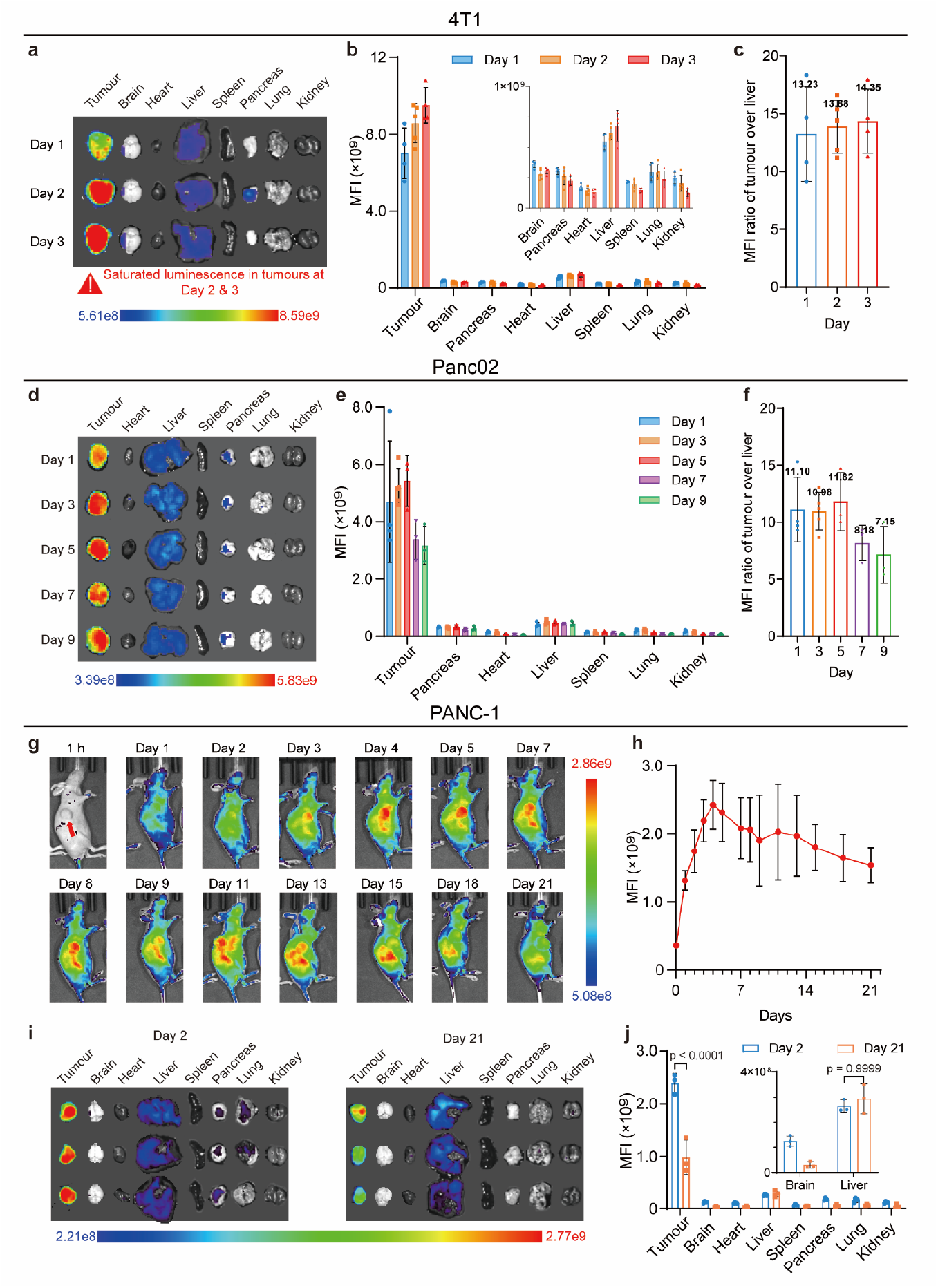
Higher dose of OEGMA-11 NPs improves accumulation and retention in the tumour. **a, b** Ex vivo fluorescence imaging (**a**) and MFI (**b**) of main organs and 4T1 tumours extracted from BALB/c mice on Day 1, 2 and 3 after tail-vein injection of NPs (40 mg/kg, n = 4 or 5). The inset in **b** shows detailed MFI of main organs. **c** MFI ratios between Panc02 tumour and the liver from **b** (R value; n = 4 or 5). **d, e** Ex vivo fluorescence imaging (**d**) and MFI (**e**) of main organs and Panc02 tumours extracted from C57BL/6 mice on Day 1, 3, 5, 7 and 9 after tail vein injection of OEGMA-11 NPs (30 mg/kg, n = 3 or 5). **f** MFI ratio between Panc02 tumour and the liver from **e** (R value). **g** Fluorescence whole-body imaging of PANC-1 tumour-bearing BALB/c nude mice for 21 days after tail vein injection of OEGMA-11 NPs (30 mg/kg, n = 3). Red arrow indicates tumour site. **h** Changes in MFI of tumours at different time points calculated based on **g. i**,**j** Ex vivo fluorescence imaging (**i**) and MFI (**j**) of main organs and PANC-1 tumours extracted from BALB/c nude mice on Day 2 and 21 after tail vein injection of OEGMA-11 NPs (30 mg/kg, n = 3). The inset in **j** shows detailed MFI of the brain and the liver. Data were presented as mean ± s.d. and statistical significance was determined using one-way ANOVA followed by Tukey’s multiple comparisons test.

We monitored in vivo biodistribution of OEGMA-11 NPs (30 mg/kg) in Panc02 tumour model over 9 days. The NPs rapidly accumulated in the tumours on Day 1 and MFI of the tumours slowly reached a maximum on Day 5 (Fig. S12). Moreover, 58.5% of maximal MFI of the tumours was still maintained on Day 9 (Fig. 3d,e). The R values reached the highest level at 11.82 on Day 5 and decreased to 7.15 on Day 9 (Fig. 3f).

A significant number of OEGMA-11 NPs remaining in the Panc02 tumours on Day 9 encouraged us to further investigate the metabolism for an even longer duration in subcutaneous PANC-1 (human pancreatic cancer cell line) tumour model in BALB/c nude mice. Fluorescence imaging of the whole mouse was performed for 21 days and MFI of the tumour region was calculated and plotted against time (Fig. 3g,h). We found that MFI at the tumour sites rapidly reached a maximum on Day 4 and declined slowly until the end of the monitoring on Day 21.

To quantitatively analyze the biodistribution, mice were sacrificed on Day 2 and 21, respectively, after injection of OEGMA-11 NPs. From ex vivo imaging, fluorescence was barely detectable from main organs except the liver on Day 21, but the tumours still exhibited obvious and dominated fluorescence (Fig. 3i,j). The R values decreased from 9.1 on Day 2 to 3.3 on Day 21, which further verified long retention in the tumour and tumour-targeting ability of OEGMA-11 NPs in subcutaneous PANC-1 tumour model. However, increasing the dose to 30 mg/kg of OEGMA-20 NPs slightly decreased the R value in PANC-1 tumour model compared to that in Panc02 tumour model (2 mg/kg). MFI of the liver from the mice injected with OEGMA-20 NPs was as high as 4.5 times of that of OEGMA-11 NPs (Fig. S13), suggesting that the dose of OEGMA-11 NPs at 30 mg/kg did not exceed the threshold of maximal uptake of the liver and their enhanced tumour accumulation by increasing the dose was not realized by overwhelming the liver^46^. These results revealed that increasing the dose to a therapeutic level significantly improves the ratio of OEGMA-11 NPs accumulating in the tumour than those in the liver, and that OEGMA-11 NPs could target three types of subcutaneous tumour models efficiently.

### Tumour-targeting ability of OEGMA-11 NPs in orthotopic PANC-1 pancreatic tumour

Motivated by the ability of OEGMA-11 NPs that universally target the subcutaneous tumours, we were curious whether OEGMA-11 NPs could be applied to orthotopic tumour, which allowed us to assess their performance in a more clinically relevant tumour microenvironment. Orthotopic PANC-1 pancreatic tumour model, as one of pancreatic ductal adenocarcinoma that is highly aggressive and fatal malignancy owing to lack of early diagnosis and poor prognosis^47^, was chosen to evaluate the performance of OEGMA-11 NPs. NPs with a dose of 30 mg/kg were administrated intravenously into tumour-bearing mice after 21, 35 and 49 days of tumour inoculation, using a group of healthy mice as control (Fig. 4a).

**Figure 4.**
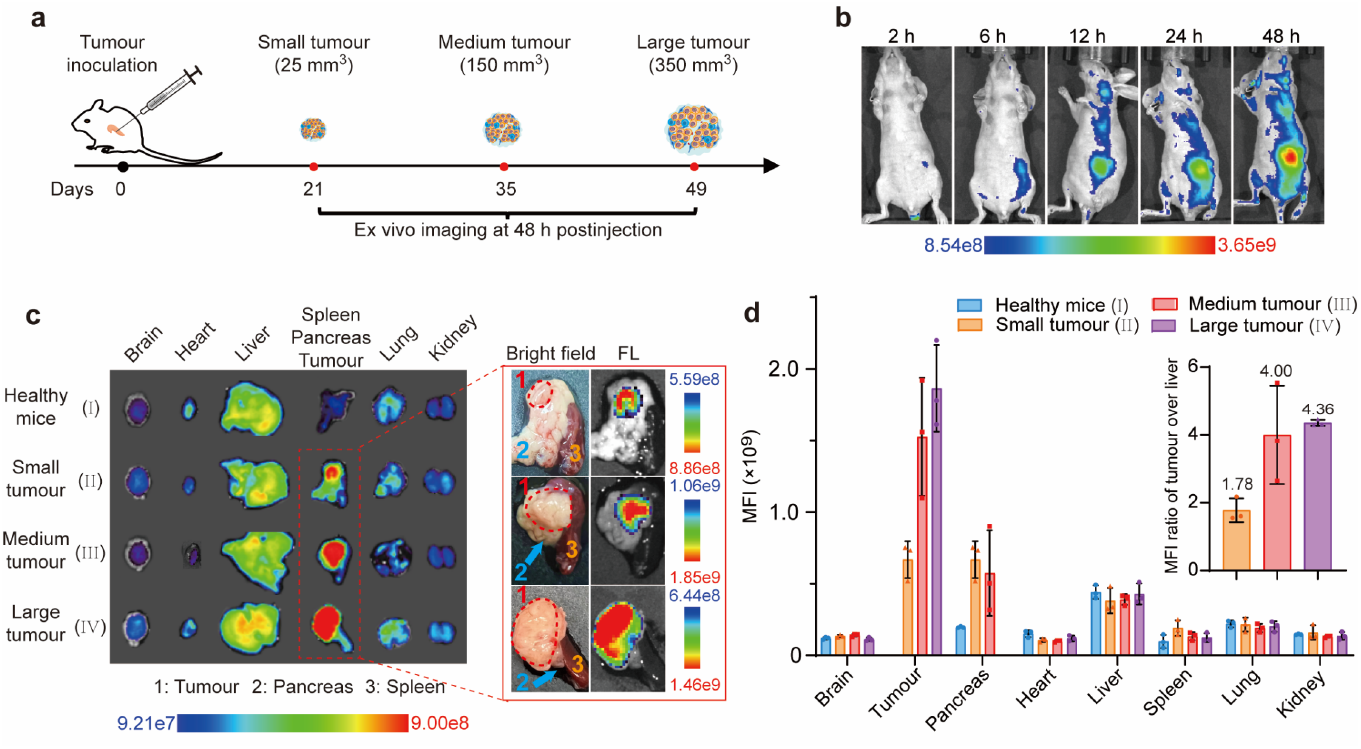
OEGMA-11 NPs exhibit tumour-targeting capabilities in orthotopic PANC-1 tumour. **a** Schematic illustration of processes of tumour inoculation, ex vivo fluorescence imaging of main organs and small, medium and large tumours at 48 h after injection of OEGMA-11 NPs (30 mg/kg). After 21, 35 and 48 days of tumour inoculation, volumes of tumour grow to 25, 150 and 350 mm3. **b** Fluorescence whole-body imaging of BALB/c nude mice bearing orthotopic PANC-1 tumour (350 mm^3^) at 2, 6, 12, 24 and 48 h after intravenous injection of OEGMA-11 NPs (30 mg/kg). **c**,**d** Ex vivo fluorescence imaging (**c**) and MFI (**d**) of main organs and PANC-1 tumours at 48 h after injection of OEGMA-11 NPs (30 mg/kg; n = 3; FL, fluorescence). MFI of the tumour and the pancreas from small tumour model in **d** was considered as equal owing to unprecise calculation of MFI of the small tumours at size of 25 mm^3^. MFI of the pancreas from the large tumour model in **d** was taken for zero owing to the large tumour occupying essentially the entire pancreas. The inset in **d** shows R value (MFI ratio of tumour over liver) of the brain and the liver. Data were presented as mean ± s.d..

As shown in Fig. 4c,d and S14, MFI of the tumour-bearing pancreas is about two folds higher than that of healthy mice injected with NPs. Apparent fluorescence signal of orthotopic pancreatic tumour with a size of 350 mm^3^ (III) was observed from the fluorescent images of the whole mice after 12 h postinjection (Fig. 4b). The tumour outline can be clearly distinguished from normal tissue of the pancreas by fluorescence imaging after 48 h postinjection, even though volumes of the tumours were as small as 25 mm^3^ (II). These results of ex vivo imaging demonstrated promising potential of OEGMA-11 NPs in the early diagnosis and treatment of pancreatic cancer. Additionally, the R values were 1.8, 4.0 and 4.4 in the small (II), medium (III) and large tumours (IV), indicating preferred accumulation in the large tumours and the efficient targeting of OEGMA-11 NPs in orthotopic PANC-1 pancreatic tumour at different stages. The relative distribution of the NPs in liver followed an order of normal mice (Fig. 5a-No.4) > mice bearing orthotopic PANC-1 tumour (Fig. 5a-No.5) > mice bearing subcutaneous PANC-1 tumour (Fig. 5a-No.7), which was the exact opposite order of accumulation of OEGMA-11 NPs in tumour (Fig. S15), implying that the tumour outcompeted with the liver in accumulation of OEGMA-11 NPs.

**Figure 5.**
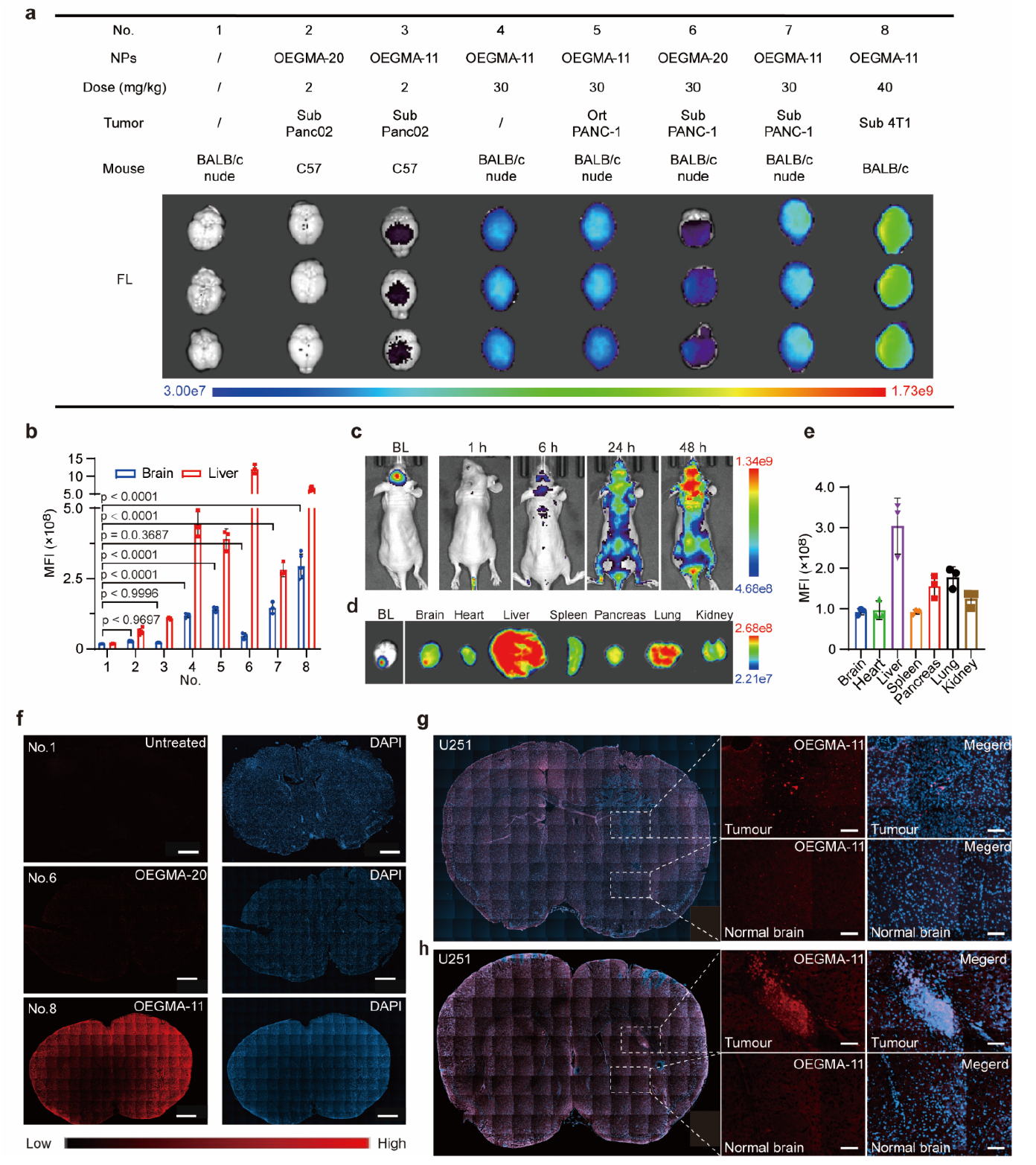
OEGMA-11 NPs can penetrate blood brain barrier and display the glioma-targeting ability. **a**,**b** Fluorescence images of the brains (**a**) and MFI of the brains and the liver (**b**) from Fig. **2b**, **3b**, **3j**, **4d** and **S13** (n = 3 or 4 or 5). Sub, subcutaneous; Ort, orthotopic. **c** In vivo bioluminescence (BL) image of orthotopic U251-luc glioma model and fluorescence whole-body imaging of mouse bearing U251-luc glioma at 1, 6, 24 and 48 h after tail-vein injection of OEGMA-11 NPs (30 mg/kg). **d**,**e** Ex vivo fluorescence imaging (**d**) and MFI (**e**) of main organs and U251-luc glioma at 48 h after tail-vein injection of OEGMA-11 NPs (30 mg/kg; n = 3). **f** Confocal images of frozen slices of the brains from (**a**) (scale bar: 1000 μm).**g**,**h** Confocal images of frozen slices of the brains from **d** (scale bar: 100 μm). Data were presented as mean ± s.d. and statistical significance was determined using one-way ANOVA followed by Tukey’s multiple comparisons test.

### Delivery of OEGMA-11 NPs to brain and glioma

We noticed the obvious fluorescence from the brain of mice during the exploration of tumour-targeting ability of OEGMA-11 NPs (Fig. 5a-b). The brain from the mice injected with OEGMA-(11, 20) NPs (Fig. 5a-No.2,3) at the dose of 2 mg/kg showed weak fluorescence comparable to the brain autofluorescence (Fig. 5a-No. 1). When the dose was increased to 30 mg/kg, nevertheless, the brains of the mice administrated with OEGMA-11 NPs showed obvious fluorescence in BALB/c nude mice (bearing tumours or not) (Fig. 5a-No.4,5,7), which was two-fold higher than that of the brains from the group of mice treated by OEGMA-20 NPs (Fig. 5a-No.6, Fig. 5b). When the dose was increased to 40 mg/kg, there was a corresponding increase of the MFI of the brains, which was 46% of MFI of the liver (Fig. 5a-No. 8) (Fig. 5b).

To further determine the entry of OEGMA-11 NPs into the brain, frozen slices of the brains were examined. As shown in Fig. 5f, with the same dose of 30 mg/kg, fluorescence of brain slices from OEGMA-11 group was much stronger than that from OEGMA-20 (Figure S13) and untreated groups, suggesting that OEGMA-11 can penetrate across intact blood brain barrier (BBB) and accumulate in the brain.

The performance of OEGMA-11 NPs inspired us to evaluate their accumulation in brain tumour model, in which U251-luc cells (human glioblastoma cell line) were used to construct orthotopic glioblastoma model. As shown in Fig. 5c, OEGMA-11 NPs gradually accumulated in the brain until 48 h. Ex vivo imaging also found that one of three mice exhibited distinct fluorescence signal in the brain tumour regions through analysis of the colocalization of fluorescence of OEGMA-11 NPs and bioluminescence (BL) of the tumour (Fig. 5d, S16). The biodistribution of OEGMA-11 NPs (30 mg/kg) in this model was basically the same as in healthy mice (Fig. 5a-No. 4, Fig. 5c,5d). We also evaluated frozen sections of the brains and found that the partial areas of the tumour exhibited enhanced fluorescence signal of OEGMA-11 NPs compared to normal brain tissues (Fig. 5f-g, S17), whereas OEGMA-7 NPs at the same dose as OEGMA-11 showed obviously less distribution in the brain and glioma (Fig. S18). Thus, these results imply that OEGMA-11 NPs are capable to infiltrate into the glioma, which demonstrates the potential applications of these NPs as carriers of imaging and/or therapeutic components targeting brain diseases.

### Antitumour efficacy and toxicity of OEGMA-11 NPs

Subcutaneous PANC-1 tumour model at the volume of about 50 mm^3^ was introduced to evaluate the antitumour effect *in vivo*. Aliquots of OEGMA-11 NPs and free CPT were intravascularly injected, respectively, into mice every 5 days at the CPT-equivalent dose of 10 mg/kg (OEGMA-11 dose = 70 mg/kg) according to literature^48^ (Fig. 3g,h and Fig. 6a). Specifically, the tumours from the mice injected with PBS as a control group grew exponentially and the mice had to be euthanized on Day 55 due to their tumour volumes over 1000 mm^3^ (Fig. 6a, b). OEGMA-11 NPs were more effective than free CPT, which gradually shrank tumours to 15 mm^3^ compared to 31 mm^3^ for free CPT-treated tumours on Day 21 after 5 doses of injections, and further delayed the tumour growth until Day 46 (Fig. 6b). The tumours in OEGMA-11 group were barely visible on Day 35 (Fig. 6d). During the whole monitoring, 86%, 70% and 43% of mice survived in OEGMA-11 NPs, free CPT and PBS groups, respectively, (Fig. 6g) and the mean weight of the existing tumours treated with OEGMA-11 NPs was 0.13 g, which was 36% of tumour weights (0.35 g) in free CPT group on Day 70 (Fig. 6b,6e,6f).

**Figure 6.**
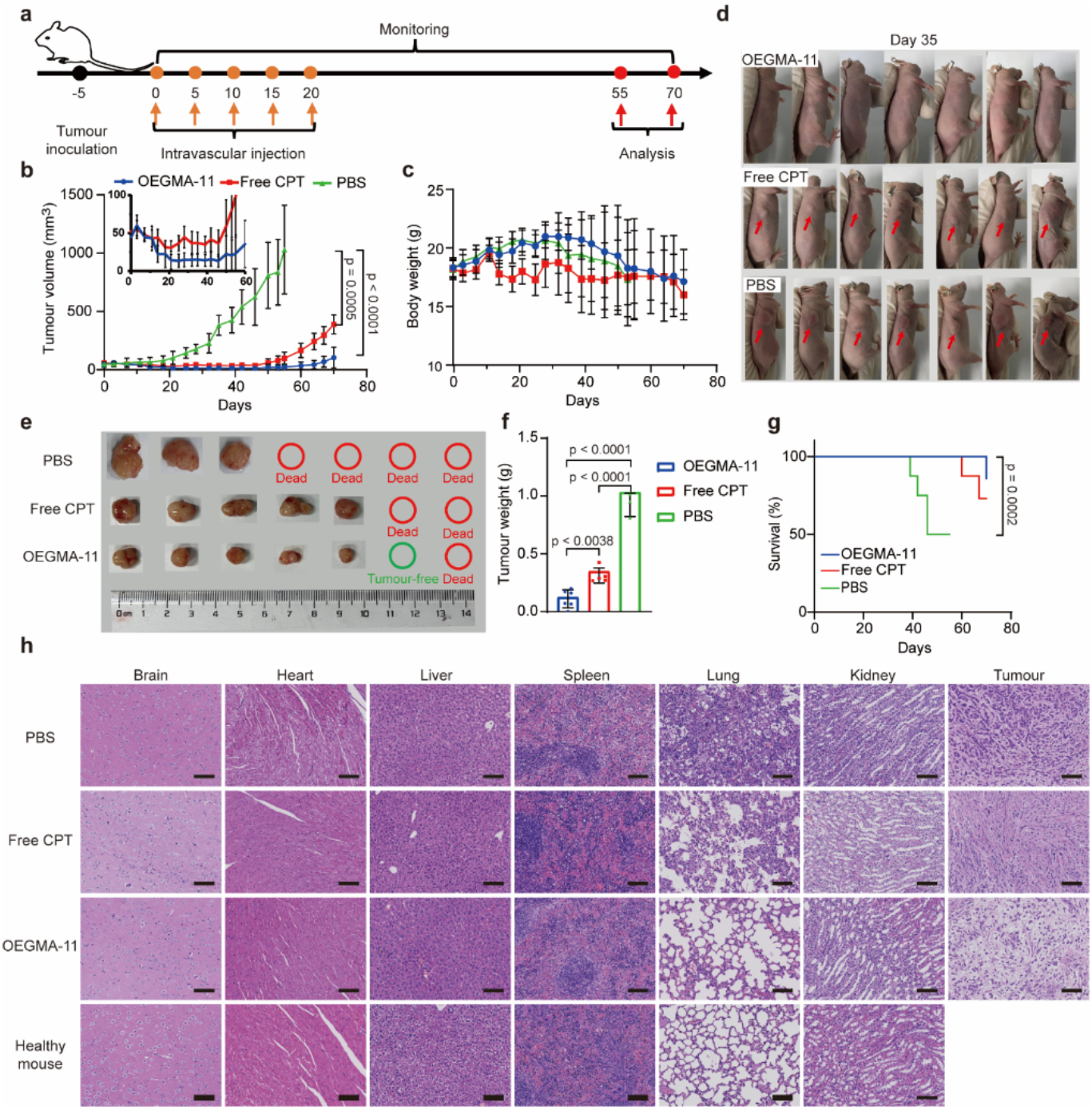
OEGMA-11 NPs show therapeutic efficacy in subcutaneous PANC-1 tumour. **a** Scheme illustration of the schedule of tumour inoculation, injeckon, monitoring of tumour size and mouse weight, analysis. **b, c** Tumour growth curves (**b**) and changes in weight (**c**) of PANC-1 bearing-mice treated with PBS, CPT (10 mg/kg) and OEGMA-11 (CPT-equivalent dose of 10 mg/kg) intravenously every five days five times (n = 7). The inset in **b** shows detailed growth curves of the mice from free CPT and OEGMA-11 groups. **d** Images of mice on Day 35 after injection of PBS, CPT and OEGMA-11. Red arrows indicate the tumour site. **e, f** Images of dissected PANC-1 tumours (**e**)and tumour weights (**f**) on 55 days (PBS group) and 70 days (free CPT and OEGMA-11 groups) (n = 3 or 5 or 6). **g** Survival curves of mice treated with PBS, CPT and OEGMA-11. **h** H&E staining images of tissues dissected from mice of PBS, free CPT and OEGMA-11 groups (Scale bar, 100 μm). Data were presented as mean ± s.d. and statistical significance in **b** was determined using one-way ANOVA followed by Tukey’s multiple comparisons test. Survival curves (**g)** were performed by the log-rank Mantel-Cox test.

In order to examine the toxicity of our OEGMA-11 NPs, hematoxylin and eosin (H&E) staining was performed and no detrimental changes were found in the brain, heart, liver, spleen and kidney of PBS, free CPT and OEGMA-11 groups. However, obvious pathological changes were observed in the lung of three groups, even in the lung of the subcutaneous tumour-free mouse from OEGMA-11 group (Fig. 6h). But the mice treated with OEGMA-11 NPs showed less pathological changes and a higher degree of apoptosis in the tumour than free CPT treatment (Fig. 6h). In addition, the mice treated with free CPT and OEGMA-11 NPs experienced 2.9 g and 0.2 g, respectively, of weight loss compared to mice in PBS group on Day 25. The body weights of mice decreased relatively slowly in the group treated by OEGMA-11 NPs compared to the group by PBS in the following 45 days (Fig. 6c).

To further evaluate the systemic toxicities after treatments, the serum chemistry analysis was conducted at the end of therapy. Serum markers, including total protein (TP), albumin (ALB), globulin (GLOB) and creatinine (CREA) of mice treated with PBS, free CPT and OEGMA-11 NPs groups were at similar levels to those of the group of healthy mice. In contrast, OEGMA-11 NPs showed some extent of interference against the level of alkaline phosphatase (ALKP) compared to the healthy mice, but the impact was more obvious compared to the group treated by PBS or free CPT (Fig. S19). Therefore, these data demonstrated enhanced therapeutic efficacy and minor adverse effect of OEGMA-11 NPs.

### The mechanism of tumour-targeting of OEGMA-11 NPs

To understand how the size and surface modification of our polymeric NPs influence their behaviors in vivo, we assessed the efficiency to surpassing the biological barriers before reaching tumour site. First, the pharmacokinetics of NPs were characterized by measuring MFI of the blood samples collected from mice at different time points. At 48 h post-injection, MFI of blood samples administered with OEGMA-11 NPs was approximately 34% of initial MFI, while negligible fluorescence was observed in blood samples of the mice administered with other NPs. OEGMA-11 NPs exhibited the longest blood-circulation half-life of about 27.0 h, compared to less than 5 h for the other NPs (Fig. 7a,b). Half-lives of MPC and CBMA NPs decreased as their sizes increased. The results of the pharmacokinetics (Table S3, Fig. 2a,b) in combination with the biodistribution as described above indicate that OEGMA-11 NPs with prolonged blood circulation time tend to escape the trap by liver and result in relatively high tumour-targeting efficacy, whereas the other NPs were more easily captured by liver, consistent with their faster blood clearance. Therefore, enhanced immune evasion could be the reason for excellent tumour-targeting of OEGMA-11 NPs.

**Figure 7.**
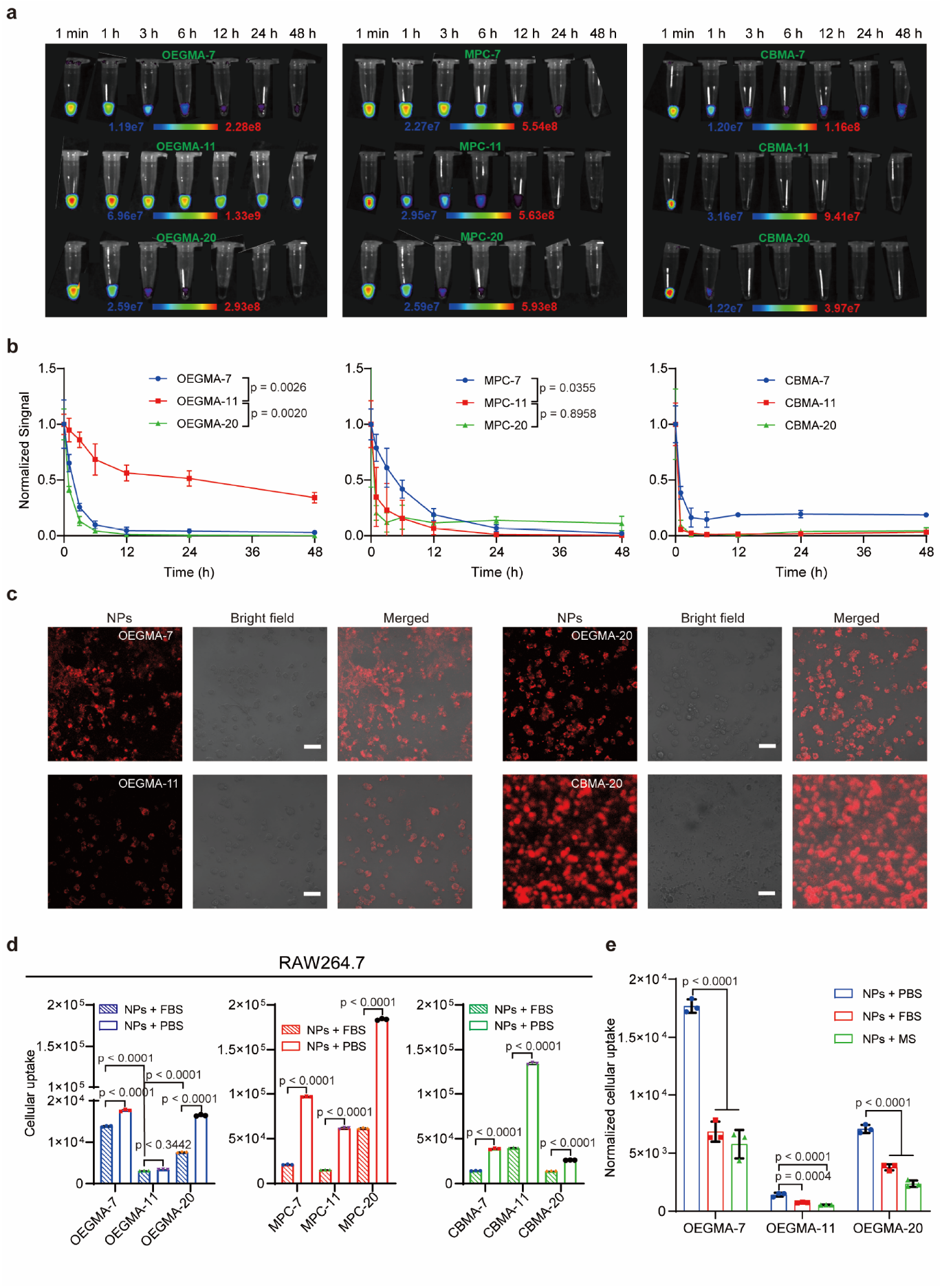
In vivo blood clearance profiles and the effect of protein corona on immune evasion of NPs. **a, b** Changes in fluorescence of blood samples collected at different time points following intravenous administration of NPs (n = 3 or 4 or 5). **c** Confocal images of RAW 264.7 cells treated with NPs after 4 h of incubation (scale bar: 100 μm). **d** Flow cytometric analysis of the cellular uptake of RAW 264.7 cells treated with NPs in serum/serum-free medium (n = 3). **e** Flow cytometric analysis of the cellular uptake of liver cells treated with FBS or mouse serum (MS) (n = 3). Data were presented as mean ± s.d.. The statistical significance in **d**,**e** was determined using one-way ANOVA followed by Tukey’s multiple comparisons test and two-tailed Student’s *t*-test were performed in **b** for comparison.

To further examine immune evasion of those NPs, the uptake by RAW 264.7 cells (mouse monocyte/macrophage-like cell) was characterized using laser scanning confocal microscope and flow cytometry. One can see that the uptake of other NPs by RAW 264.7 cells was 1.5∼19.3-fold higher than that of OEGMA-11 NPs (Fig. 7c,d). The trend of the cellular uptake of these NPs by macrophages is essentially opposite compared to the trend of their blood-circulation lifetimes as discussed above (Table S3).

Recent studies, for example, by Siegwart’s and Xu’s groups^49-52^, have shown that the physicochemical properties of NPs significantly affect the formation of protein corona in blood and that protein corona could mediate organ-specific delivery in lipid NPs. In view of the large difference in zeta potential and size between our polymeric NPs (Fig. 1e), we hypothesized that the protein corona on the surface of NPs may regulate the uptake by macrophages, the blood circulation time and the accumulation of NPs in liver. To examine this hypothesis, firstly, the formation of protein corona on the surface of our polymeric NPs was proven by sodium dodecyl sulphate-polyacrylamide gel electrophoresis (SDS-PAGE) after incubation with mouse serum (Fig. S23). Secondly, the polymeric NPs were incubated with RAW 264.7 cells in the medium with/without the presence of Fetal Bovine Serum (FBS) to control the formation of protein corona. Little change in the cellular uptake of OEGMA-11 NPs was observed in the present/absence of FBS, while other NPs experienced 25-200% increase of uptake in the FBS-free environment compared to that in the FBS environment (Fig. 7d). These results suggest that the formation of protein corona decreased the uptake of the NPs by macrophages.

We further used liver cells collected from the liver of C57BL/6 mice to mimic the clearance in vivo and further explore the effect of protein corona. The OEGMA-7, -11 and -20 NPs pretreated with PBS, FBS or mouse serum (MS) were incubated with liver cells in serum-free medium. The results (Fig. 7e) show that the uptake of OEGMA-11 NPs remained minimal under these three conditions and the internalization of all NPs by liver cells followed an order of PBS > FBS > MS, which is consistent with the cellular uptake by RAW 264.7.

Subsequently, the effect of protein corona on the uptake in cancer cell line was also investigated. Specifically, there was an increase of the internalization of all NPs by PANC-1 cells in serum-free medium compared to that in the cell-incubation media in the presence of FBS (Fig. S24).

The results above suggest that the outstanding pharmacokinetics and tumour-targeting capability of OEGMA-11 NPs, determined by a synergistic effect of their physicochemical properties (size, surface charge, hydrophobic/hydrophilic ratio, rigidity and hairy surface feature), could be related to their unique performance in avoidance of the sequestering by mononuclear phagocytic systems such as macrophages existing in blood, liver, spleen.

To further determine the transcellular transport of NPs through blood vessels of tumour, bEnd.3 cells (mouse brain endothelial cell line) were used to culture in the upper chamber separated by a microporous polycarbonate membrane to mimic tumour vasculature (Fig. 8b). Then NPs of OEGMA-7, 11, 20 and MPC-7, due to their relatively long circulation time, were added to the medium in the upper chamber and the capability of transcellular transport could be defined by measuring the fluorescence intensity of the medium in the basal chamber. The MFI of basal medium from OEGMA-11 group was significantly higher compared to that treated by the other NPs during the whole monitoring, which was about 6.4-, 5.2- and 2.1-fold higher than that of OEGMA-7, 20 and MPC-7 NPs at 48 h, respectively, indicating high transcellular transport efficiency of OEGMA-11 NPs (Fig. 8c) which also accounts for their outstanding penetration into the brain tissues and tumours.

**Figure 8.**
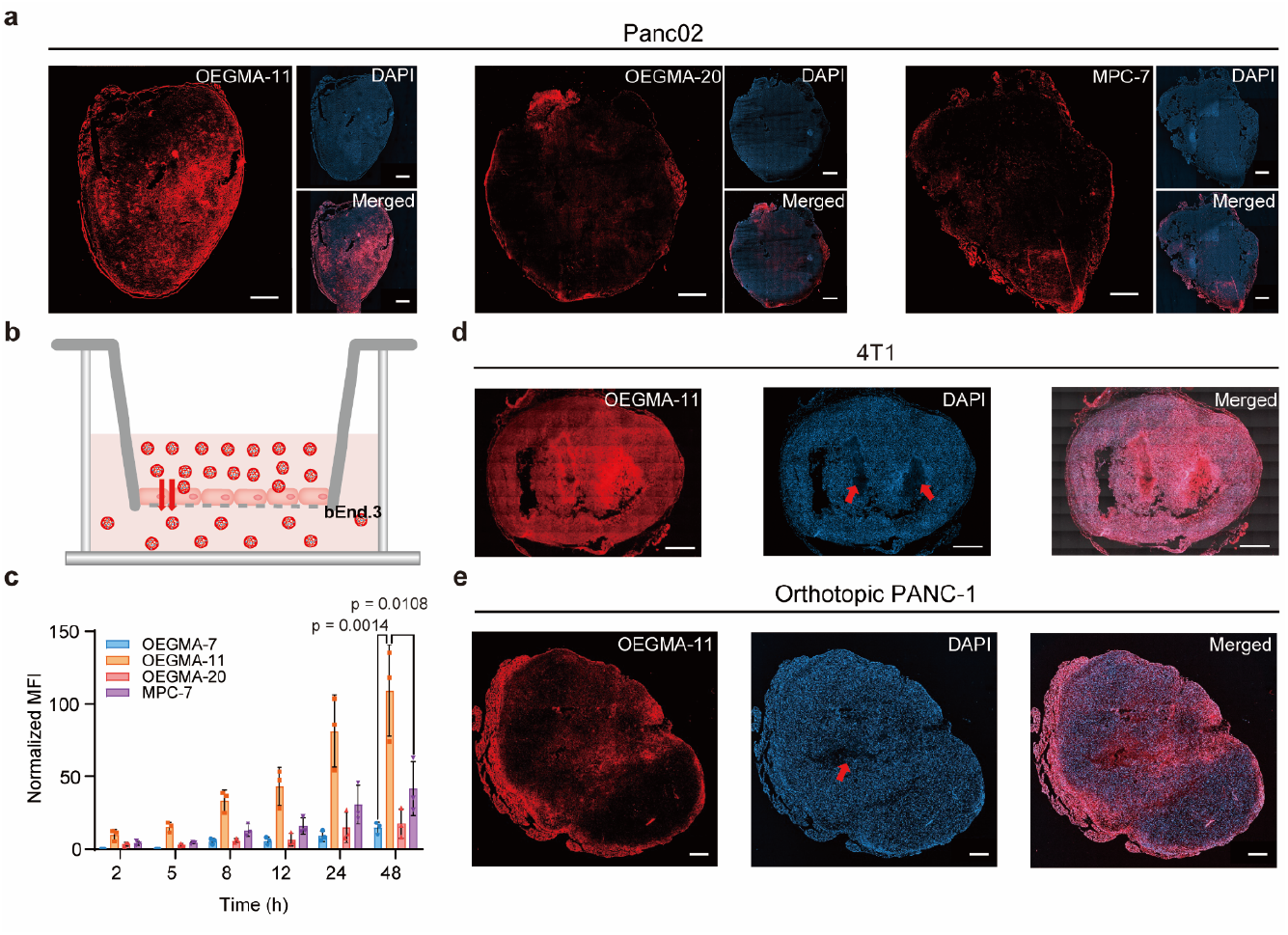
Transcytosis and penetration of NPs. **a** The tumour penetration of NPs in Panc02 tumour at 48 h postinjection (Scare bar: 1000 μm). **b** Schematic illustration of transwell model for investigating the transcellular transport of NPs. **c** MFI of the medium in lower chamber at different time points after adding the NPs (n = 3). **d**,**e** The tumour penetration of OEGMA-11 NPs in 4T1 tumour (**d**) and orthotopic PANC-1 tumour at 48 h postinjection (Red arrows: necrosis; Scare bar: 1000 μm). Data were presented as mean ± s.d and statistical significance was determined using one-way ANOVA followed by Tukey’s multiple comparisons test.

Next, we analyzed the tumour penetration by fluorescence-imaging the distribution of NPs in the cryo-sectioned tumours. Fig. 8a shows that most regions of tumour slice from the mice treated with OEGMA-11 NPs exhibited strong fluorescence even at the center of Panc02 tumour slice with a volume of about 200 mm^3^. Similar deep tumour-penetration depth of OEGMA-11 NPs was also observed in the slices of aggressive 4T1 tumour model (Fig. 8d and S10b). In contrast, the fluorescence images of the tumour slices from the mice treated with OEGMA-20 or MPC-7 NPs showed that the NPs mainly accumulate in the outer peripherical region of the tumour slice, but barely in the central region of the tumour. However, OEGMA-11 NPs primarily distributed in outer periphery and necrosis region in slices of orthotopic PANC-1 tumour with a volume of about 350 mm^3^, compared to the penetration of the same NPs in subcutaneous 4T1 and Panc02 tumour models (Fig. 8e). In addition, the release of CPT from OEGMA-11 NPs was observed in the Panc02 tumour slice through analysis of the colocalization of fluorescence of CPT (blue) and the polymer (red) (Fig. S21).

To evaluate the tumour-targeting capability of our polymeric NPs in vitro, we measured the uptake of these NPs in a series of cancer cell lines (4T1, PANC-1 and Panc02) using flow cytometry. As shown in Fig. S24-26, the cellular uptake of OEGMA-11 NPs by these three cancer cell lines is the lowest among all the tested NPs, similar to the cellular uptake by the macrophages as shown in Fig. 7d. The uptake of CBMA NPs, due to their positive surface zeta potential, by cancer cells was the highest among all the tested NPs, which was over 100-fold higher than that of OEGMA NPs. The uptake of MPC NPs by PANC-1 cells was similar to that of OEGMA NPs, while 4T1 and Panc02 cells internalized about 1 to 10 times more MPC NPs than OEGMA NPs (Fig. S24-26).

Overall, the results above show that OEGMA-11 NPs with superior immune evasion avoided the fast clearance in blood circulation, resulting in prolonged blood-circulation lifetime, which provided more opportunities for the continuous entry to the tumour and brain and maintained the long-term retention of the NPs in tumours. In contrast, the other bottle-brush polymeric NPs, including MPC- and CBMA-series, intravascularly administrated in mice were rapidly internalized by macrophages and liver cells and then cleared at a relatively slow rate upon the formation of protein corona, finally trapped most inside the liver (for MPC polymers) and the lung (for CBMA polymers). In addition, the high transcellular transport through tumour vasculature and deep tumour penetration of OEGMA-11 NPs may further improve the delivery efficiency to the tumour tissues.

## Discussion

We have presented a series of theranostic bottle-brush polymers with bright fluorescence and high stability in water and revealed the relationship between the size and the surface chemistry of our polymeric NPs and their selective targeting behaviors against tumour, liver, brain and lung both in vivo and in vitro levels. Remarkably, the universality of passive tumour-targeting capability of OEGMA-11 NPs has been demonstrated in three subcutaneous tumour models, an orthotopic pancreatic tumour model and an orthotopic glioblastoma model implanted in three types of mice, suggesting their promising clinical applications in diagnosis and treatment of various cancers. We have also studied the tumour-targeting mechanism of OEGMA-11 NPs from the following aspects: reduced clearance of immune cells, prolonged blood circulation time, facilitated extravasation through tumour vasculature and deep infiltration into tumours. Moreover, the formation of protein corona may protect this type of polymeric NPs from the quick clearance by MPS and impede their uptake by cancer cells. Our results may offer insight for the rational design of polymeric delivery systems with optimal structures and physicochemical properties to overcome cascaded biological barriers and improve the targeting efficacy against tumours and other inflammation diseases.

## Methods

### Materials

11-Bromo-1-undecanol, 3,4-dihydro-2H-pyran, p-toluenesulfonic acid monoandryte, 2,7-dibromofluorene, 4,7-Dibro-2,1,3-benzothididiazole, tris(dibenzylideneacetone)dipalladium (Pd_2_(dba)_3_), tris(2-methoxyphenyl) phosphine ((o-MeOPh)_3_P), pivalic acid (PivOH), 3-hexylthiophene, Camptothecin (CPT), MgSO_4_, oligo(ethylene glycol) methyl ether methacrylate (OEGMA), azobisisobutyronitrile (AIBN), 2-methacryloyloxyethyl phosphorylcholine (MPC), 2-Hydroxyethyldisulfide, hydroquinone, triethylamine, acrylylchloride, camptothecin (CPT), Dimethylaminopyridine (DMAP), Dicyclohexylcarbodiimide (DCC), triphosgene, 2-(Dimethylamino) ethyl methacylate, tert-butylbromoacetate, KOH, KI, K_2_CO_3_ were purchased from Sigma-Aldrich (China) or Macklin (China) or Aladdin (China). Dulbecco’s modified Eagle’s medium (DMEM), RPMI-1640 medium, fetal bovine serum (FBS), penicillin-streptomycin, Matrigel matrix and phosphate buffered saline (PBS) were purchased from Gibco (USA).

### Preparation and characterization of polymeric NPs

A proper amount of OEGMA, MPC and CBMA polymers were dissolved in organic solvent (THF for OEGMA polymers; ethanol and THF (volume ratio:1:1) for MPC polymers; DMF and THF (volume ratio:1:1) for CBMA polymers) and the polymer solution was rapidly injected into water to obtain dispersion of polymeric NPs. The dispersion was transferred into a dialysis bag with a MWCO of 7 kDa (ThermoFisher, USA) for dialysis against deionized water to remove organic solution and stored at 4 C° for further use. Surface zeta potential and hydrodynamic size of polymeric micelles were measured by Malvern 3000HS Zetasizer (Malvern Instruments Ltd, UK). Morphologies of polymeric micelles were observed by transmission electron microscopy (TEM, Gatan, USA) at 80 KV. All NPs for biological experiments were diluted in 1×PBS buffer.

### Establishment of mouse tumour model

The care and use of laboratory animals were performed according to the approved protocols of the Institutional Animal Care and Use Committee (IACUC) at The Chinese University of Hong Kong, Shenzhen and Shenzhen Institute of Translational, Shenzhen Second People’s Hospital. The tumour volume was calculated by the formula: V = 1/2 × (tumour length) × (tumour width)^2^.

To establish subcutaneous 4T1 tumour model, 1×10^7^ 4T1 cells in 150 μL of 1×PBS were injected in the right flank region of the mice (BALB/c, female, 4-5 weeks old, Charles River, USA).

To establish subcutaneous PANC-1 tumour model, 1×10^7^ PANC-1 cells in 150 μL of 1×PBS were injected in the right flank region of the mice (BALB/c nude, female, 4-5 weeks old, Charles River, USA).

To establish subcutaneous Panc02 tumour model, 1×10^7^ Panc02 cells in 150 μL of 1×PBS were injected in the right flank region of the mice (C57BL/6, female, 4-5 weeks old, Charles River, USA).

To establish orthotopic PANC-1 tumour model, mice (BALB/c nude, female, 4-5 weeks old, Charles River, USA) were anesthetized by isoflurane and general area of the spleen was located. Using surgical scissor makes an incision and 3×10^5^ PANC-1 cells mixed with matrix gel in 20 μL of 1×PBS were injected into the tail of pancreas. Then the incision was closed by suture.

To establish glioblastoma model, mice (BALB/c nude, female, 4-5 weeks old) were anesthetized by isoflurane and fixed on a stereotactic frame (RWD, Shenzhen, China). 3×10^5^ U251-luc cells in 3 μL of 1×PBS were stereotactically injected into the parenchyma of brain (brightlateral: 2.0 mm, bregma: 1.8 mm, depth: 3 mm).

### Cellular uptake

The PANC-1, 4T1, Panc02 and RAW 264.7 cells were used to examine the cellular uptake of the polymeric micelles. The three types of cells at a density of 5×10^5^ cells per well were seeded in 6-well plates and cultured overnight. The medium containing 100 μg/mL polymeric micelles or NPs was incubated with cells for 4 h. The cells were washed with 1× PBS, digested by tyrisin, washed again with 1× PBS and the cellular uptake was measured by flow cytometry (Cytek, DxP Athena, USA).

For the analysis of uptake of liver cells, liver cells were obtained by grinding liver (from C57BL/6 mouse) through cell strainer (40 μm). 100 μg/mL NPs were premixed with PBS/FBS/MS (mouse serum) and added into cell medium (FBS-free) for 4 h incubation. The cells were washed with 1× PBS, digested by tyrisin, washed again with 1× PBS and the cellular uptake was measured by flow cytometry or confocal microscopy (LSM800, Zeiss, Germany).

### Transcytosis

The b.End 3 cells were used to examine the ability that NPs passed through blood vessels in vitro. The cells at a density of 5×10^5^ cells per well were seeded in transwell (0.4 μm, Corning, USA) and cultured for 3 days to cover the whole bottom of well. The medium containing 100 μg/mL NPs were added into the upper chamber and normal medium were added into the lower chamber. The medium from the lower chamber was measured for fluorescence intensity using microplate reader (Syneryy H1, BioTek, USA) at 2, 5, 8, 12, 24 and 48 h.

### Pharmacokinetic study

200 μL NPs at the dose of 2 μg/g were injected into tail vein of C57BL/6 mice and 40 μL blood was collected at 0, 1, 3, 6, 12, 24 and 48 h, which was mixed with 4 μL EDTA (15 mg/mL). The plasma was obtained by centrifugation at 500 g for 5 min and measured for MFI using IVIS (PerkinElmer, USA).

### Biodistribution studies of polymeric NPs

200 μL NPs at the dose of 2 μg/g were injected into tail vein of mice bearing subcutaneous Panc02 or 4T1 tumours (n = 5 or 3, tumour volume of ∼ 200 mm^3^). Mice were euthanized at 48 h after administration and major organs (Brain, heart, liver, spleen, lung and kidney) and tumours were harvested. MFI of those tissues was measured by IVIS (Excitation: 520 nm; Emission: 660 nm).

To determine the effect of dose on biodistribution and metabolism of NPs, 200 μL NPs at the dose of 40 μg/g were injected into tail vein of mice bearing subcutaneous 4T1 tumours (n = 4, tumour volume of ∼ 250 mm^3^). Mice were euthanized on Day 1, 2 and 3 after administration and major organs (Brain, heart, pancreas, liver, spleen, lung and kidney) and tumours were harvested. MFI of those tissues was calculated by IVIS. For mice bearing subcutaneous Panc02 tumours (n = 4, tumour volume of ∼ 200 mm^3^), 200 μL NPs at the dose of 30 μg/g were injected into tail vein of mice. Mice were euthanized on Day 1, 3, 5, 7 and 9 after administration and major organs (heart, pancreas, liver, spleen, lung and kidney) and tumours were harvested. MFI of those tissues was calculated by IVIS. For mice bearing subcutaneous PANC-1 tumours (n = 3, tumour volume of ∼ 200 mm^3^), 200 μL NPs at the dose of 30 μg/g were injected into tail vein of mice. Mice were imaged for 21 days and euthanized on Day 2 and 21 after administration. Major organs (brain, heart, pancreas, liver, spleen, lung and kidney) and tumours were harvested and MFI of those tissues was calculated by IVIS.

To study the biodistribution in orthotopic PANC-1 tumour model, 200 μL NPs at the dose of 30 mg/kg were injected into tail vein of mice (n = 3) after 21 or 35 or 49 days of tumour inoculation. Mice were euthanized at 48 h after administration and major organs (Brain, heart, pancreas, liver, spleen, lung and kidney) and tumours were harvested. MFI of those tissues was calculated by IVIS.

To study the biodistribution in U251-luc orthotopic tumour model, 200 μL NPs at the dose of 30 mg/kg were injected into tail vein of mice (n = 3). Mice were euthanized at 48 h after administration and major organs (Brain, heart, pancreas, liver, spleen, lung and kidney) were harvested. MFI of those tissues was calculated by IVIS.

### In vivo tumour and brain penetration

The tumour and organ tissues were obtained from the experiments of biodistribution of NPs. The tumours and organs were immersed in 4% paraformaldehyde (PFA) solution, sucrose solution, paraffin, respectively, and were cut into slices at the thickness of 10 μm in a cryostat. The slices were observed by confocal microscopy (LSM800, Zeiss, Germany).

### Antitumour therapy in PANC-1 subcutaneous model

The mice at 50 mm^3^ of tumour volume were randomly divided into three groups (n = 7). 5 doses of PBS, free CPT (10 mg/kg) and OEGMA-11 NPs (10 mg/kg CPT-equivalent dose) were injected via tail vein every five days and the tumour volume and mouse weight were monitored for 70 days (PBS group for 55 days). At the end of the experiment, organs and serum from mice were collected for further evaluation of cytotoxicity.

### In vivo cytotoxicity

For serum biochemistry analysis, serum was obtained from the experiment of antitumour therapy and total protein (TP), albumin (ALB), globulin (GLOB), creatinine (CREA) and alkaline phosphatase (ALKP) were measured using biomedical analyzer (Catalyst One, IDEXX, USA).

For histological analysis, slices of organ tissues form the experiment of tumour therapy were obtained by performing the same procedures of frozen slices, a stained with hematoxylin and eosin and imaged by microscope (ImageXpress Pico, SQS-40P, China).

## Acknowledgements

We thank the financial support by the University Development Fund-Research Start-up Fund (UDF01001806) from the Chinese University of Hong Kong, Shenzhen, and Shenzhen Science and Technology Innovation Commission (JCYJ20220818102804009). We thank the Advanced Materials Laboratory and the School of Science and Engineering for access to instrumental platform such as ^1^HNMR and TEM. W.Z. thanks the financial support of Ph.D. Scholarship from the Chinese University of Hong Kong, Shenzhen.

## Author contributions

M.W. conceived the concept and directed the research. M.W., H.T., W.Z. and Y.X. designed the experiments. W.Z. synthesized and characterized the polymers. Y.X. constructed the tumour models. W.Z., and Y.X. performed the biological experiments. R.G. contributed to the instrumental usage of animal experiments. P.Z., and H.H. helped the biological experiments. W.Z. and M.W. wrote the manuscript with inputs from all coauthors. All authors read and commented on the manuscript.

