## Supporting Information for "Theranostic Bottle-Brush Polymers Tailored for Universal Solid-Tumour Targeting"

**Synthetic routes**

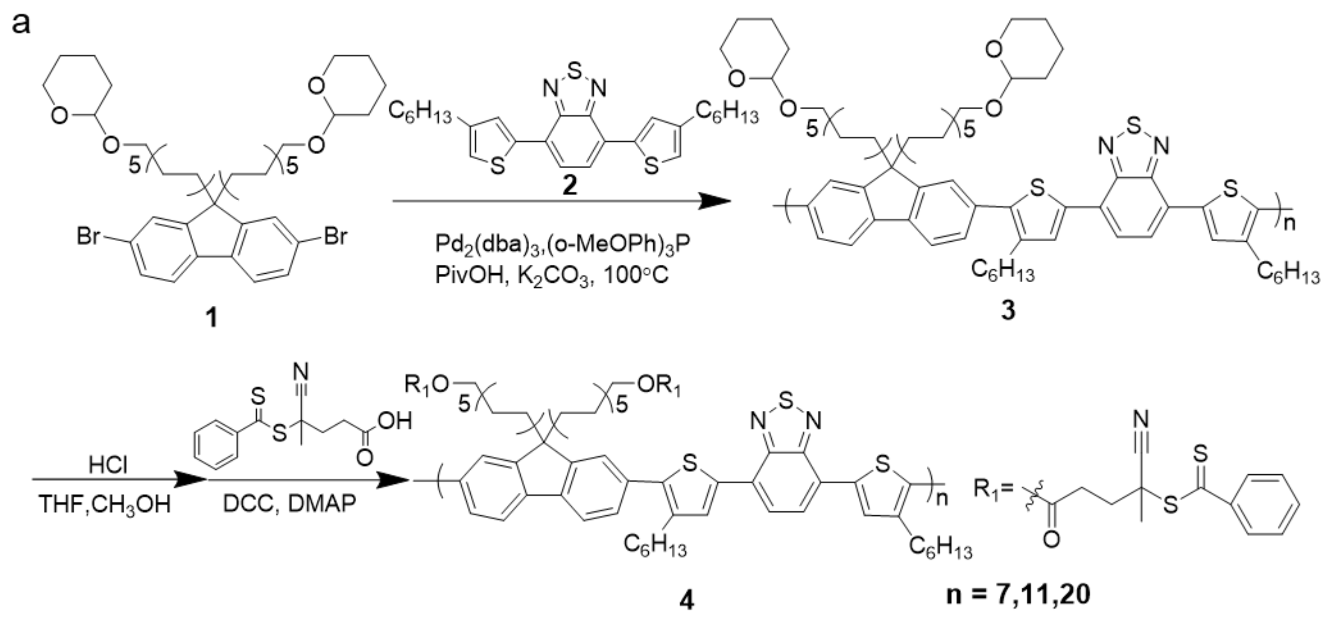


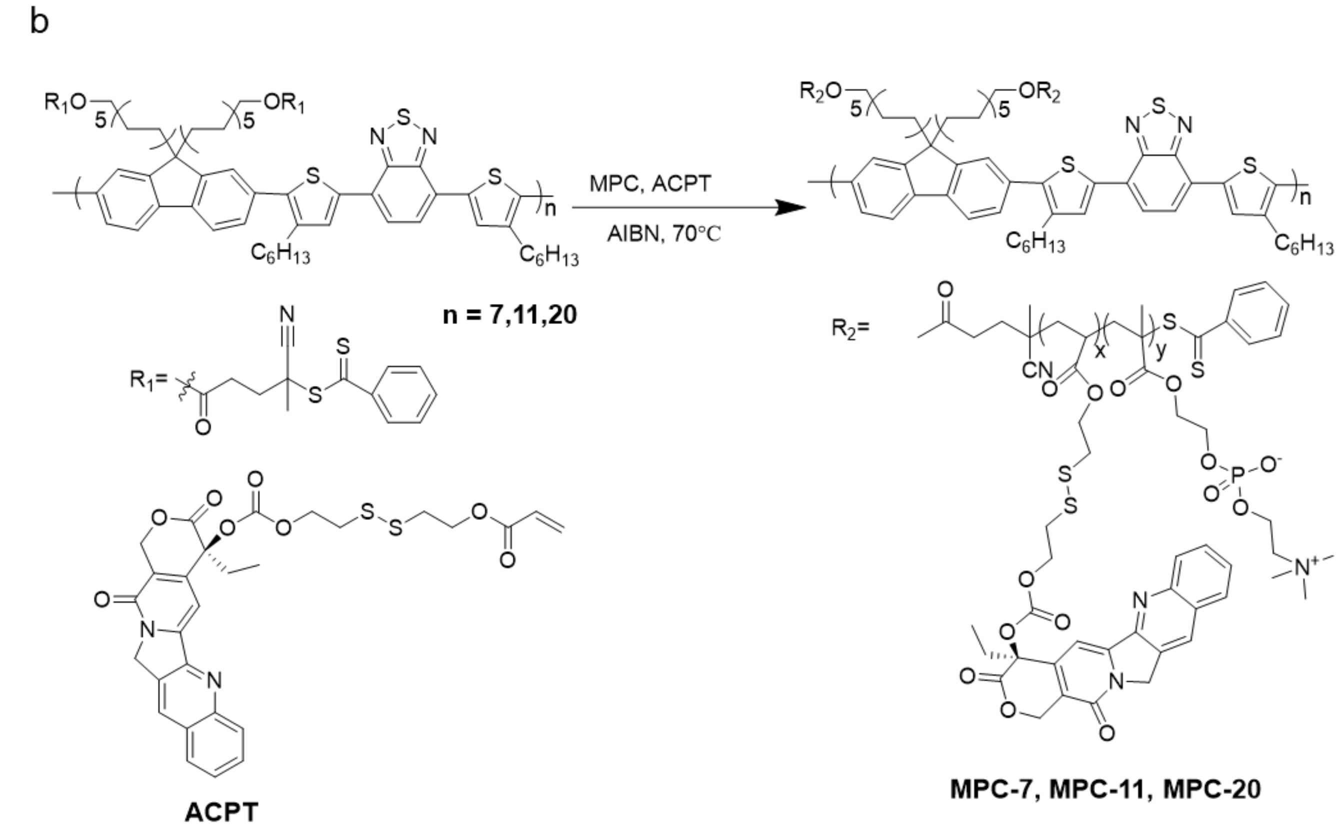


**Scheme S1**: Synthesis route to polymers 4 (a) and MPC polymers (b).

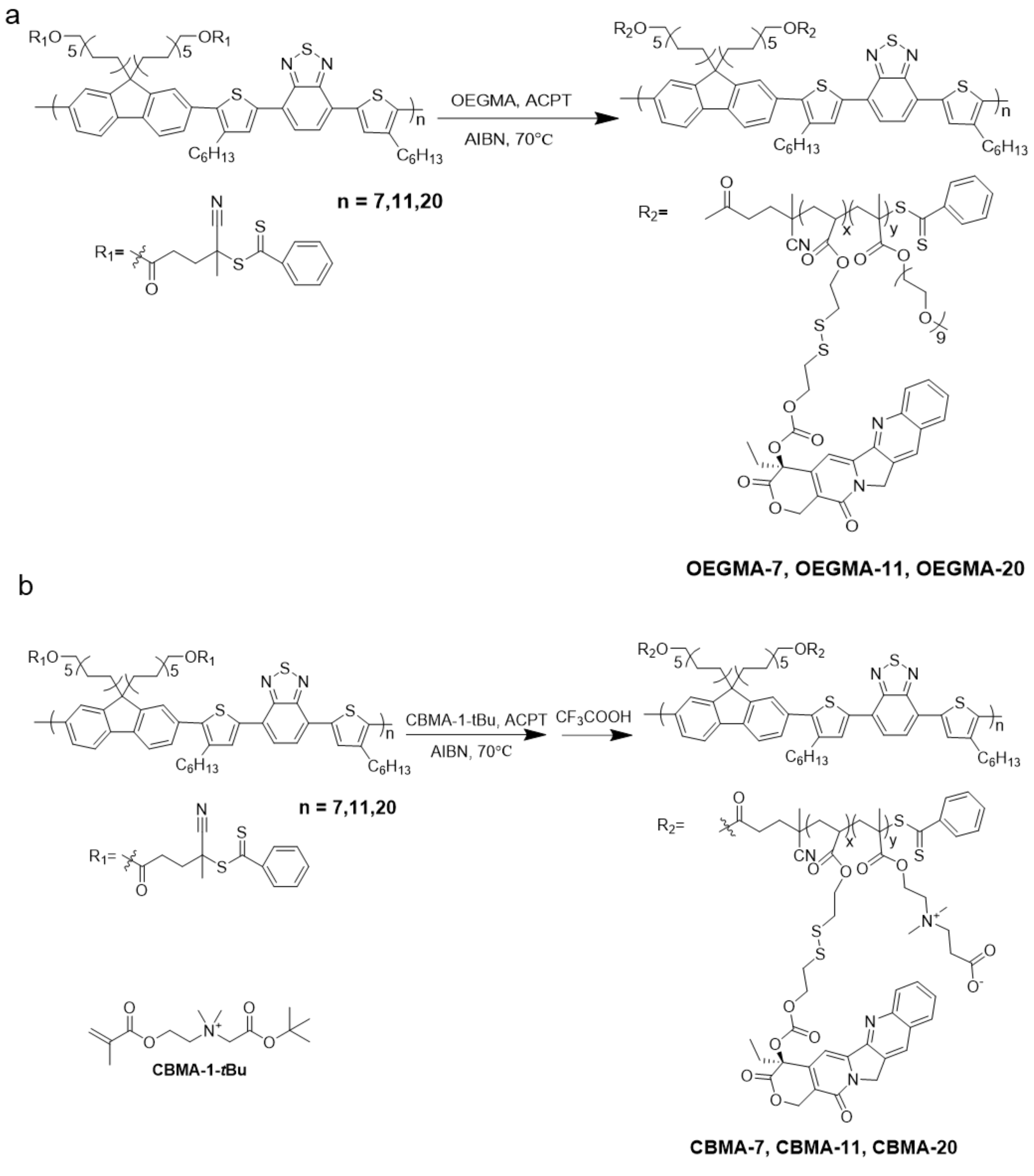


**Scheme S2**: Synthesis route to OEGMA polymers (a) and CBMA polymers (b).

**Synthesis of ACPT, CBMA-1-*t*Bu, monomer 1, 2, 3** can be found in literatures[1-3].

**Synthesis of polymer 4:** Under a $N_2$ atmosphere, polymer 3 (120 mg) pretreated by HCl, 4-Cyano-4-(phenylcarbonothioylthio) pentanoic acid (RAFT agent, 0.492 mmol, 137.45 mg), DCM (30 mL) and 4-(Dimethylamino) pyridine (DMAP) (0.0246 mmol, 3 mg) were added and the solution was placed into ice bath. Dicyclohexylcarbodiimide (DCC, 1.5 mmol, 0.344 mL) in 5 mL dry DCM was added via a micro syringe. The temperature of the flask was slowly raised to room temperature and the reaction was continued for 24 h. The solution was precipitated and filtered in cold methanol[3].

**Synthesis of OEGMA polymers:** Polymer 4 (20 mg), oligo(ethylene glycol) methyl ether methacrylate (OEGMA, 0.465 mmol, 232.5 mg) and CPTA monomer (0.093 mmol, 54.2 mg) were dissolved in 1 mL of THF in a Schlenk tube. And azobisisobutyronitrile (AIBN) in THF solution (1 mL × 0.01 mol/L) was added to the mixture. After five freeze-and-thaw cycles, the tube was then heated at 70 °C for 48 h. A solution of the crude product was precipitated in cold diethyl ether for three times[4].

**Synthesis of CBMA polymers:** Polymer 4 (20 mg), CBMA-1-tBu (0.465 mmol, 126.5 mg), AIBN (1.6 mg) and CPTA monomer (0.093 mmol, 54.2 mg) were dissolved in 4 mL of solution (DMF : chloroform = 1:1) in a Schlenk tube. After five freeze-and-thaw cycles, the tube was then heated at 70°C for 72 h. A solution of the crude product was precipitated in cold diethyl ether for three times. Product (100 mg) in 4 mL solution (DMF:Chloroform=1:1) was mixed with 1 mL trifluoroacetic acid and solution was stirred for 10 h. A solution of the crude product was precipitated in cold diethyl ether for three times.

**Synthesis of MPC polymers (Scheme 2.4): Polymer 4** (20 mg), 2-methacryloyloxyethyl phosphorylcholine (MPC, 0.465 mmol, 137.3 mg), AIBN (1.6 mg) and CPTA monomer (0.093 mmol, 54.2 mg) were dissolved in 4 mL of solution (ethanol : chloroform = 1:1) in a Schlenk tube. After five freeze-and-thaw cycles, the tube was then heated at 70°C for 72 h. A solution of the crude product was precipitated in cold diethyl ether for three times.

**Cell culture:** Human pancreatic cancer cell line (PANC-1), mouse pancreatic epithelial cell line (Panc02) and mouse brain endothelial cell line (b. End 3) were cultured in DMEM, Murine macrophage-like cell (RAW 264.7) and mouse breast cancer cell line were cultured in RPMI-1640 medium. All the medium were added with 10% FBS and 1% penicillin-streptomycin. All the cells are purchased from American Type Culture Collection (ATCC) and cultured at 37 °C with 5% $CO_2$.

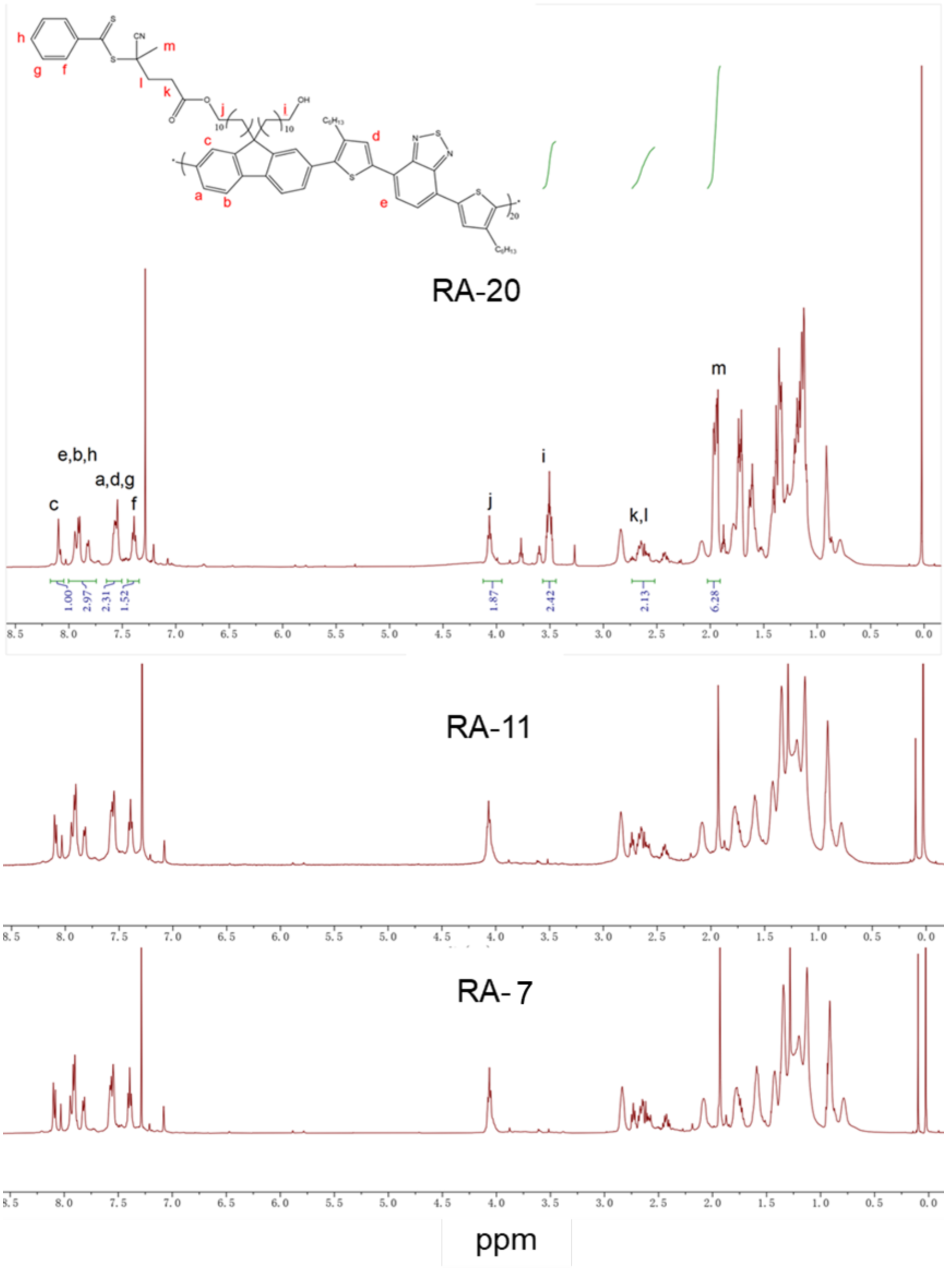


**Figure S1.** $^{1}$HNMR (500 MHz) spectra of polymer 4 in $CDCl_3$ at 25 °C.

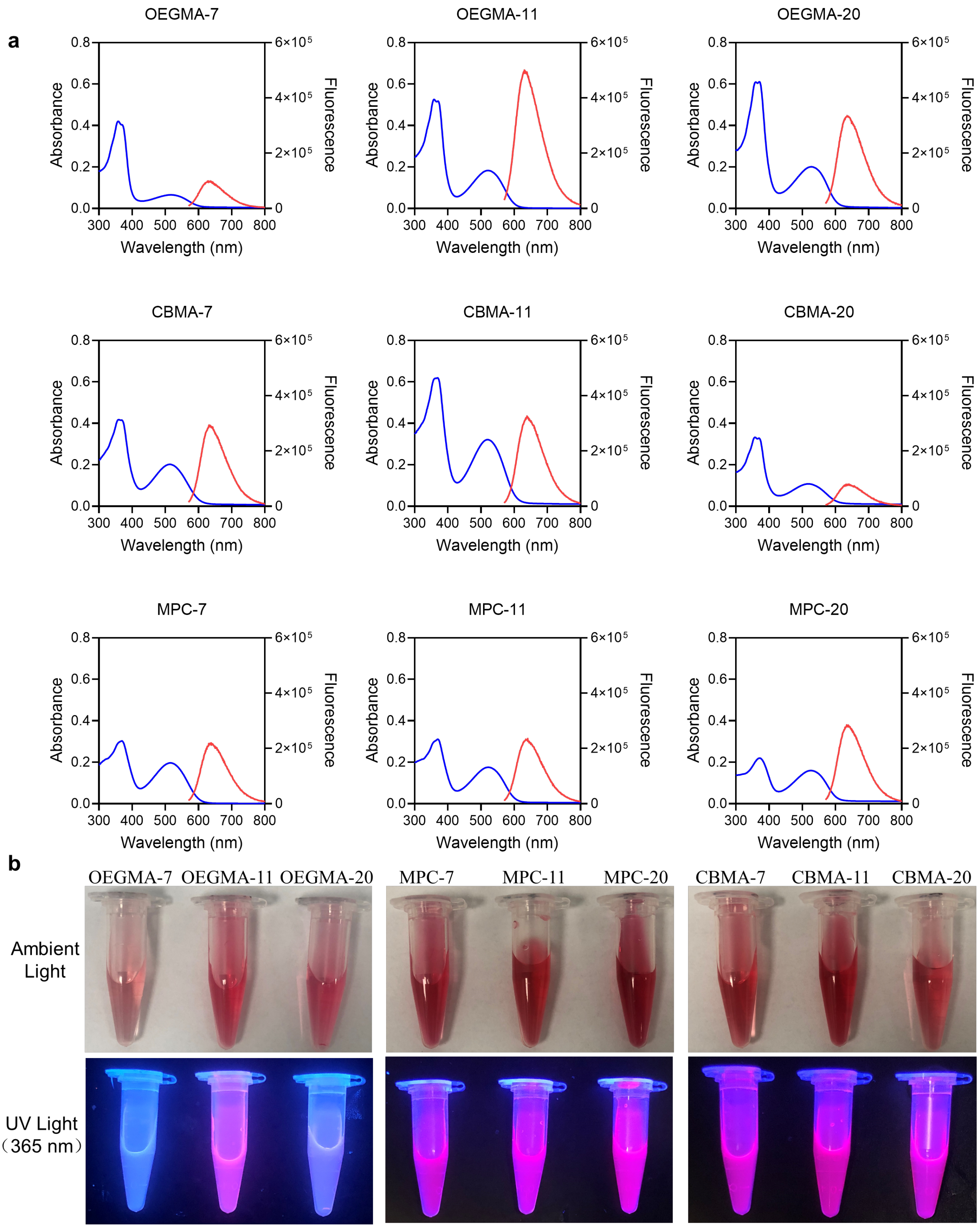


**Figure S2. a** The UV-Vis absorption and steady-state fluorescence emission spectra (excitation: 520 nm) of NPs in water; **b** Digital photographs of NPs under ambient lignt and UV light (365 nm).

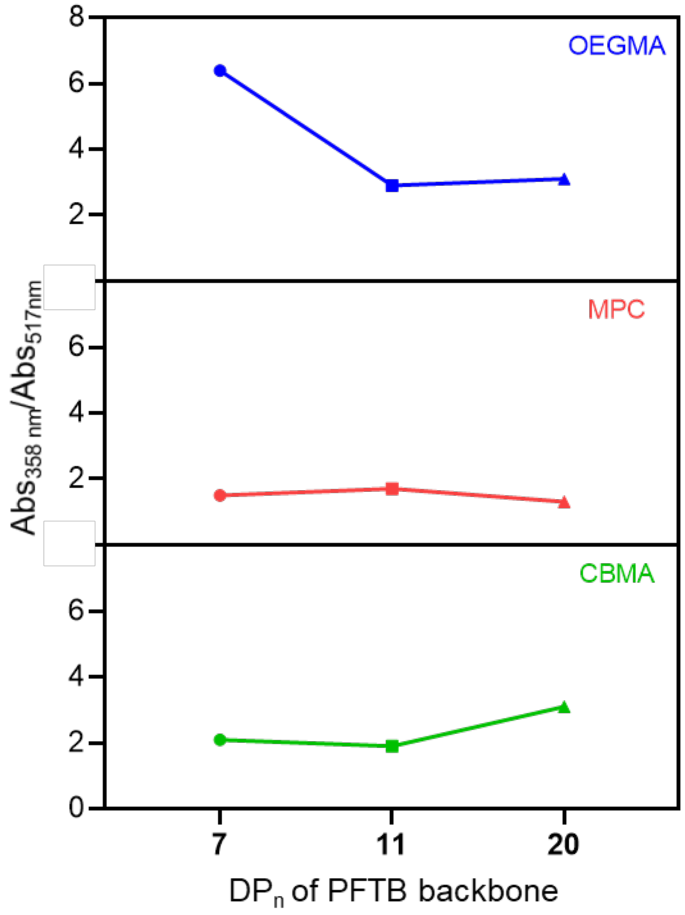


**Figure S3.** UV-vis absorbance ratio of CPT (358 nm) over PFTB (517 nm) in different polymers ($DP_n$: number-average degree of polymerization).

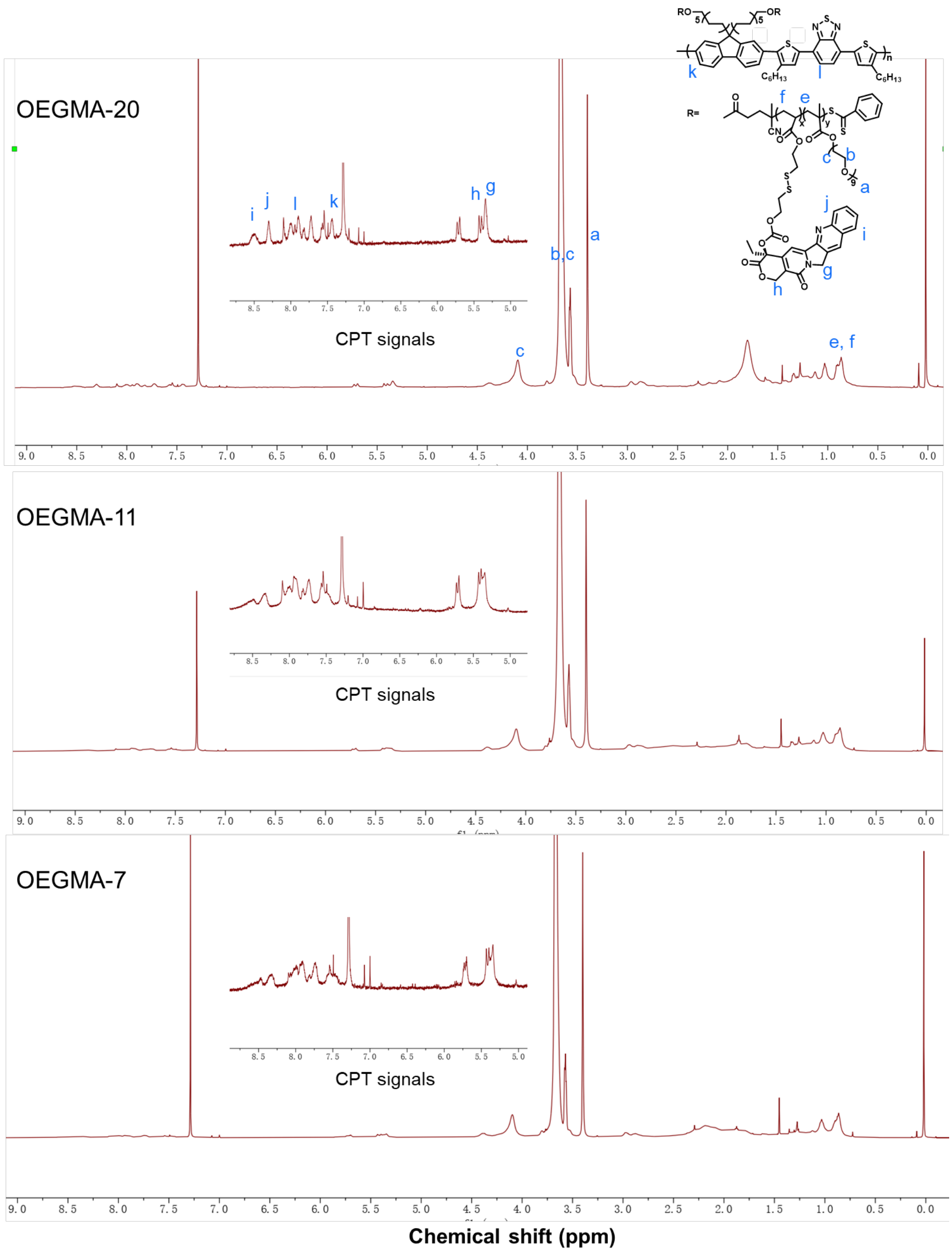


**Figure S4.** $^{1}$HNMR (500 MHz) spectra of OEGMA polymers in $CDCl_3$ at 25 °C.

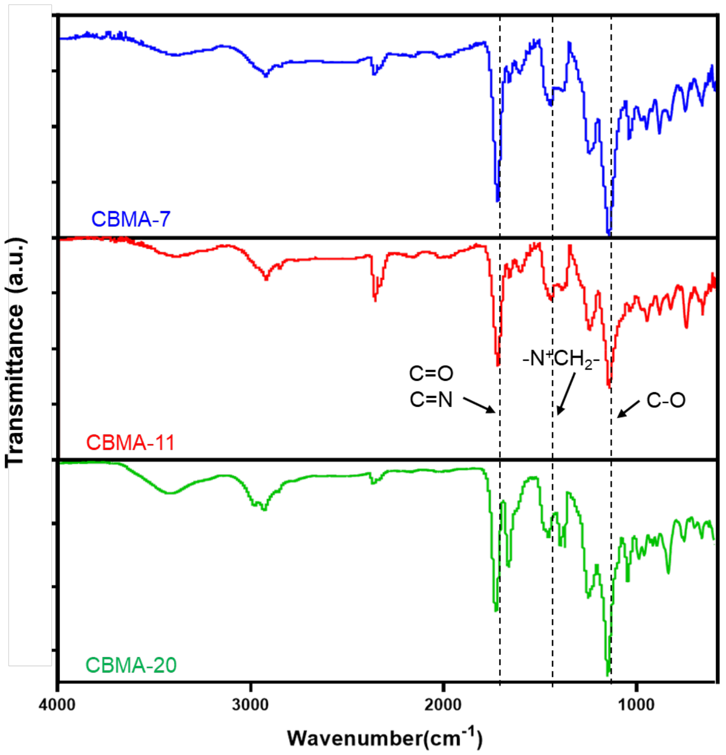


**Figure S5.** FTIR spectra of CBMA polymers.

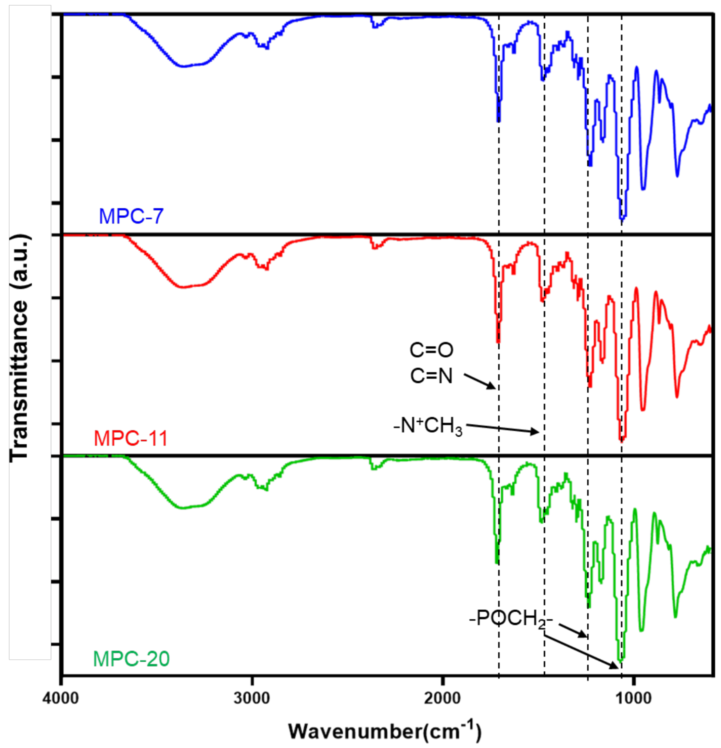


**Figure S6.** FTIR spectra of MPC polymers.

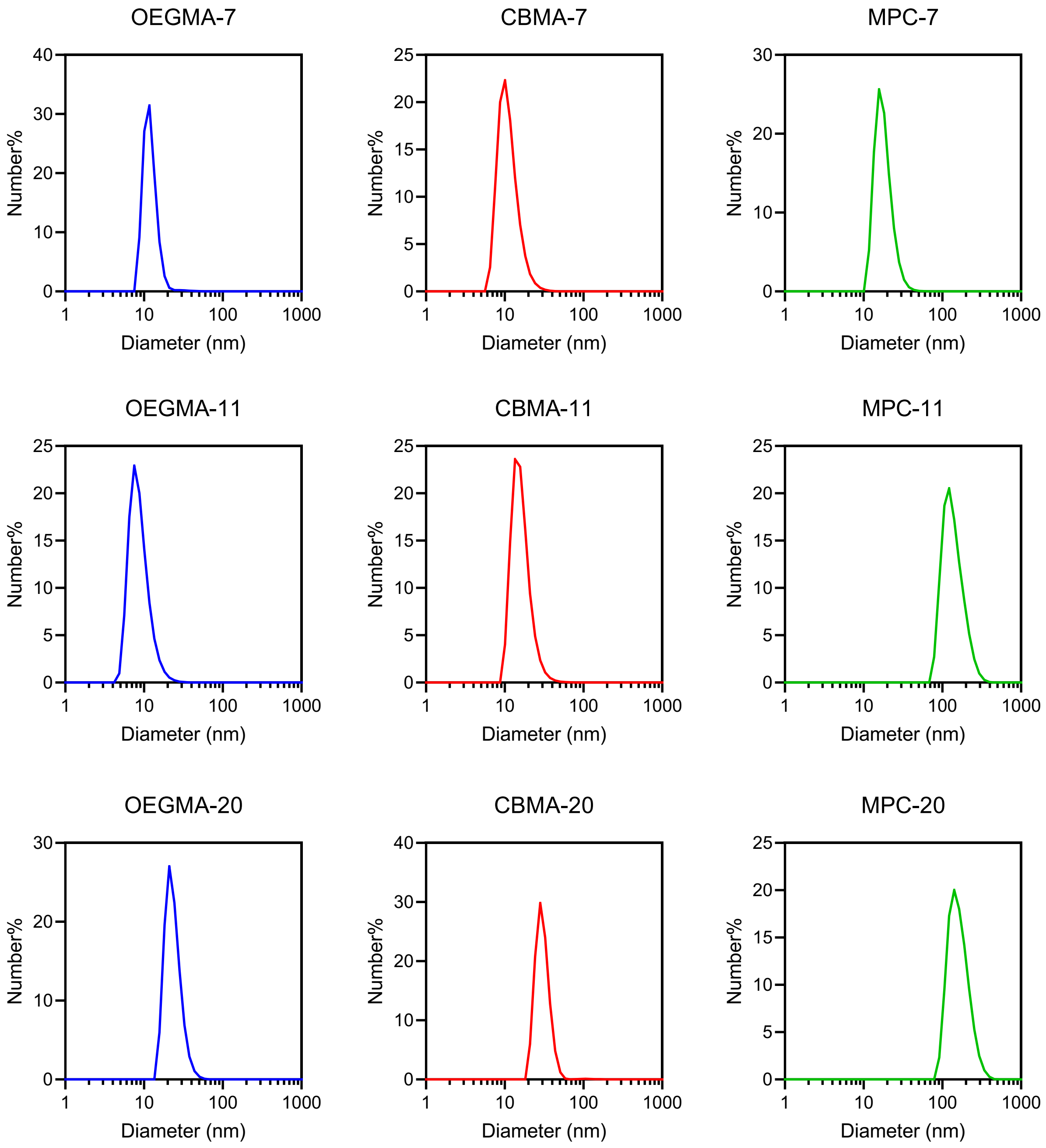


**Figure S7.** DLS results of OEGMA-7,11,20 NPs, CBMA-7,11,20 NPs and MPC-7,11,20 NPs in water.

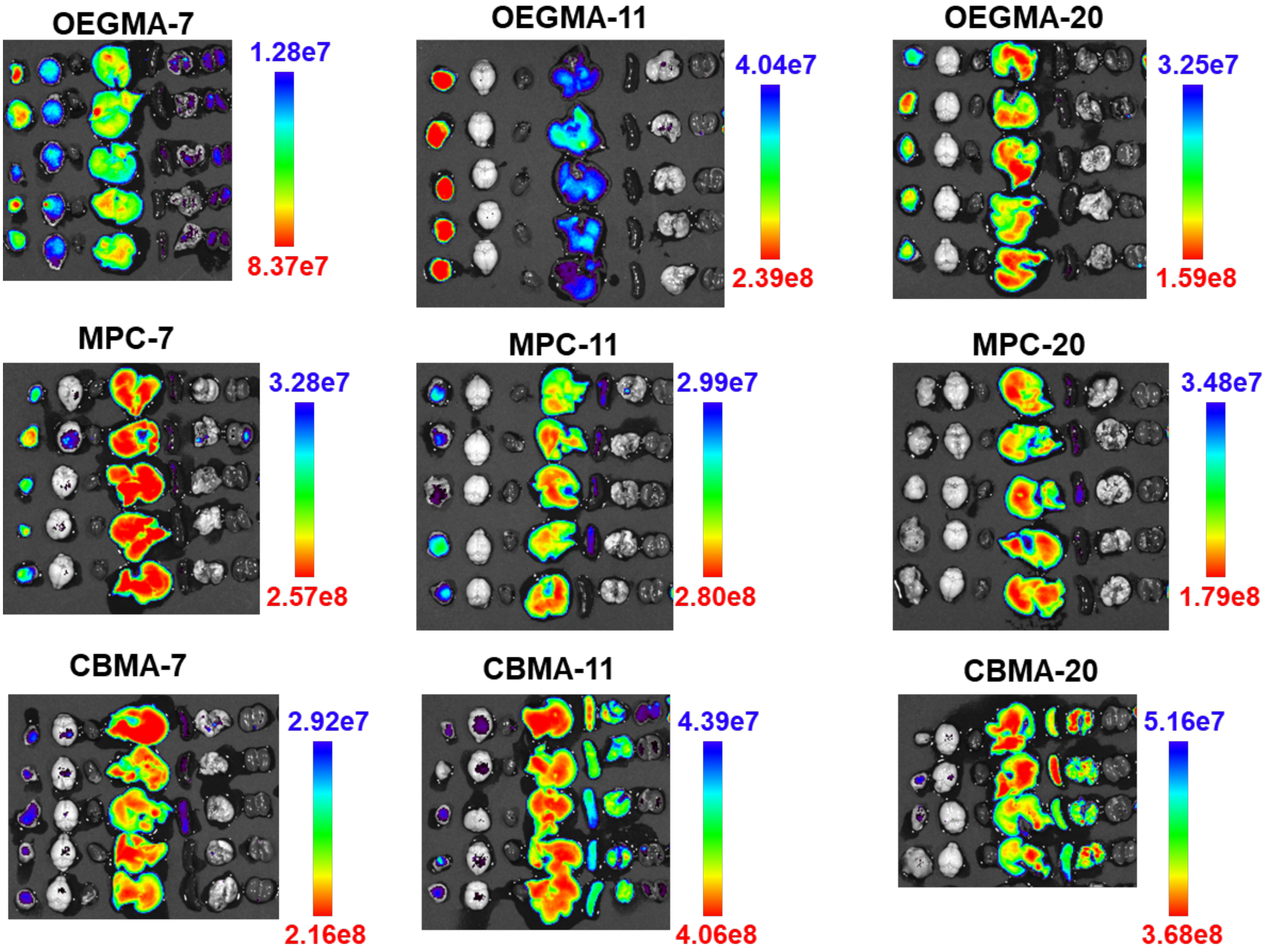


**Figure S8.** Ex vivo fluorescence imaging of major organs in C57BL/6 mice bearing Panc02 subcutaneous tumour at 48 h administration (organs from left to right: tumour, brain, heart, liver, spleen, lung, kidney; Dose of NPs: 2 mg/kg).

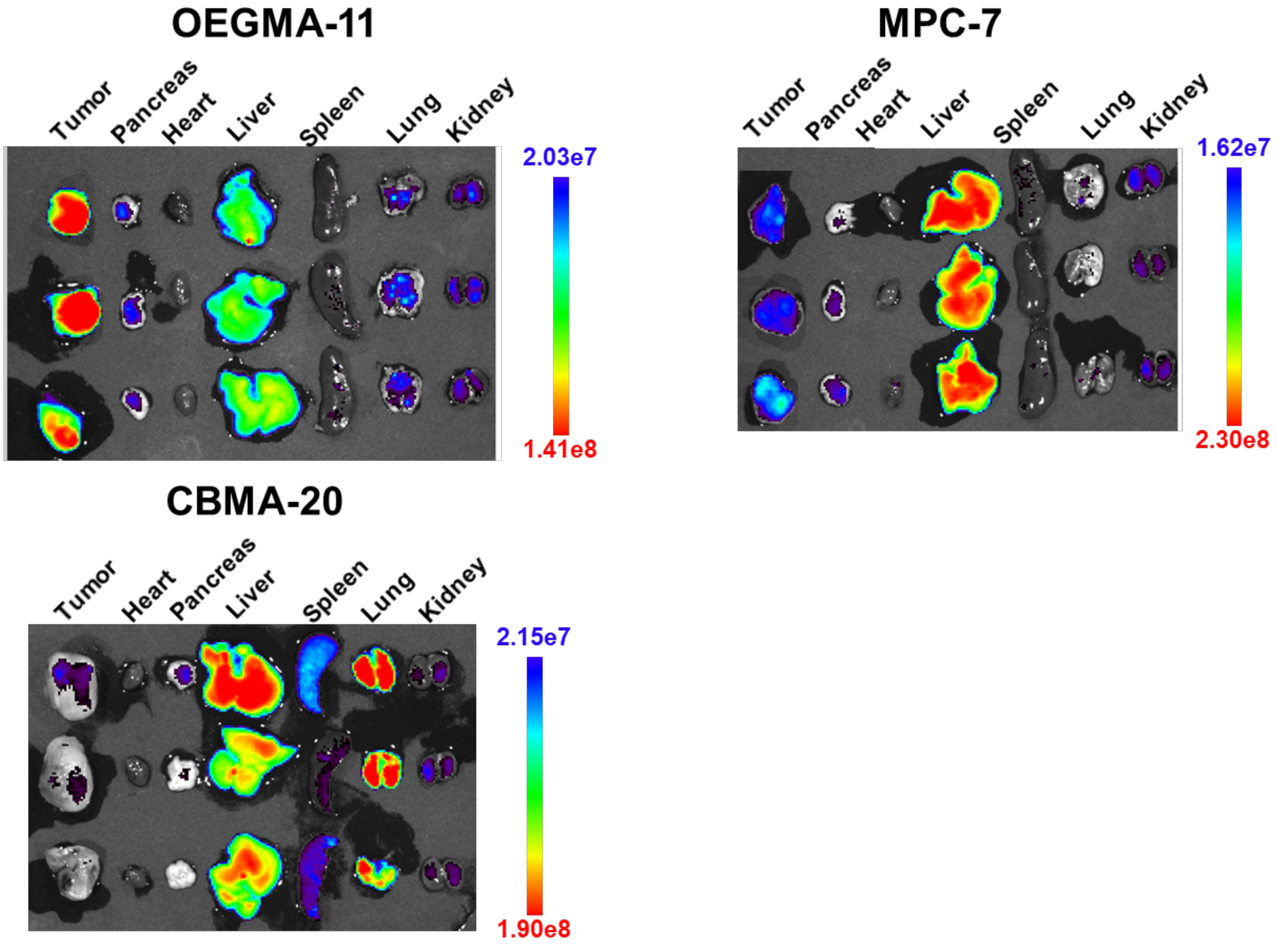


**Figure S9.** Ex vivo fluorescence imaging of major organs in BALB/c mice bearing 4T1 subcutaneous tumour at 48 h administration (Dose of NPs: 2 mg/kg).

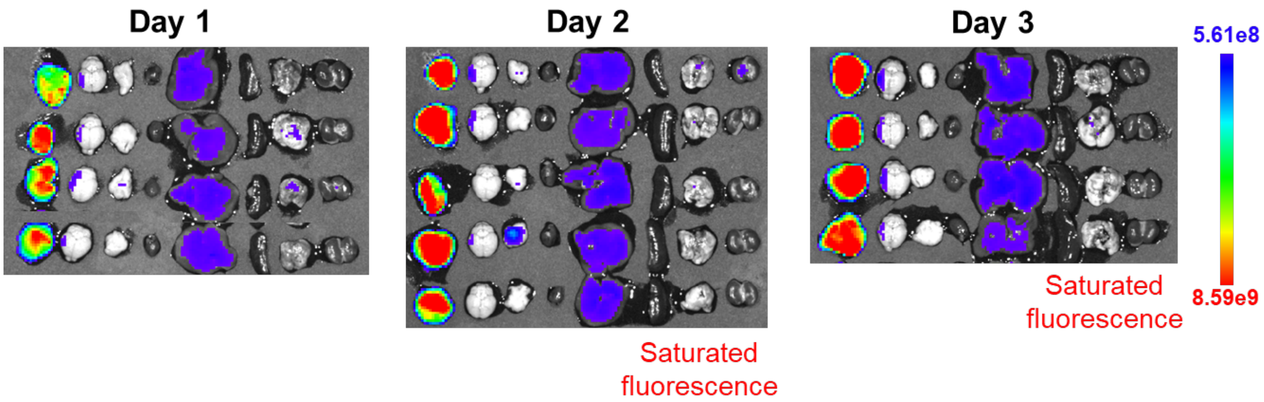


**Figure S10.** Ex vivo fluorescence imaging of major organs in BALB/c mice bearing 4T1 subcutaneous tumour after 1, 2, 3 days postinjection with OEGMA-11 NPs (organs from left to right: tumour, brain, pancreas, heart, liver, spleen, lung, kidney; Dose of NPs: 40 mg/kg).

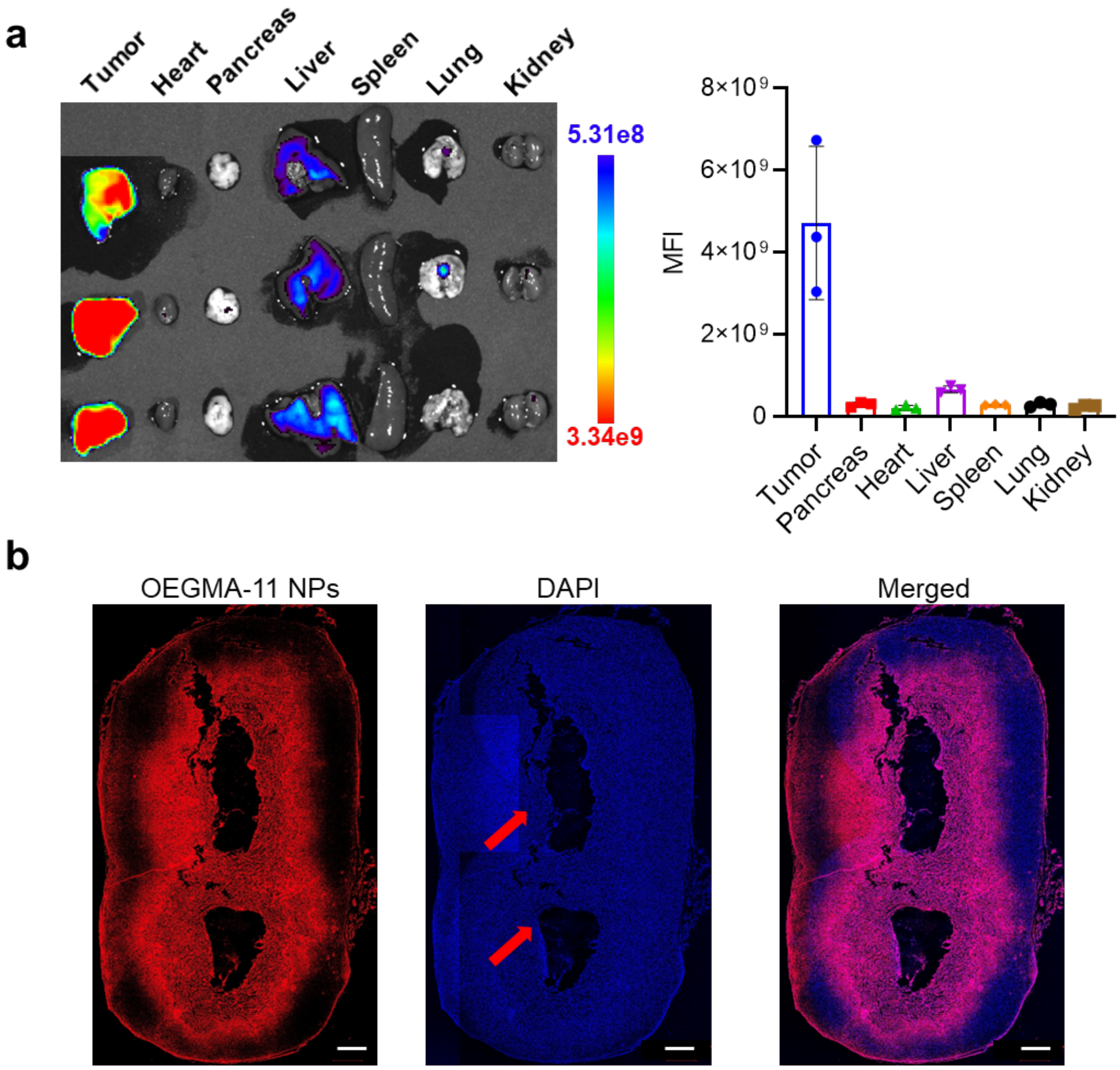


**Figure S11.** The biodistribution and penetration of OEGMA-11 NPs (Dose of NPs: 30 mg/kg). **a** Ex vivo fluorescence imaging and MFI of major organs and 4T1 tumours (with necrosis) after 48 h administration of OEGMA-11 NPs. **b** Fluorescence images of cryo-slice of 4T1 tumour (with necrosis) after 48 h administration of OEGMA-11 NPs (scale bar: 1000 μm, red arrow indicates necrosis).

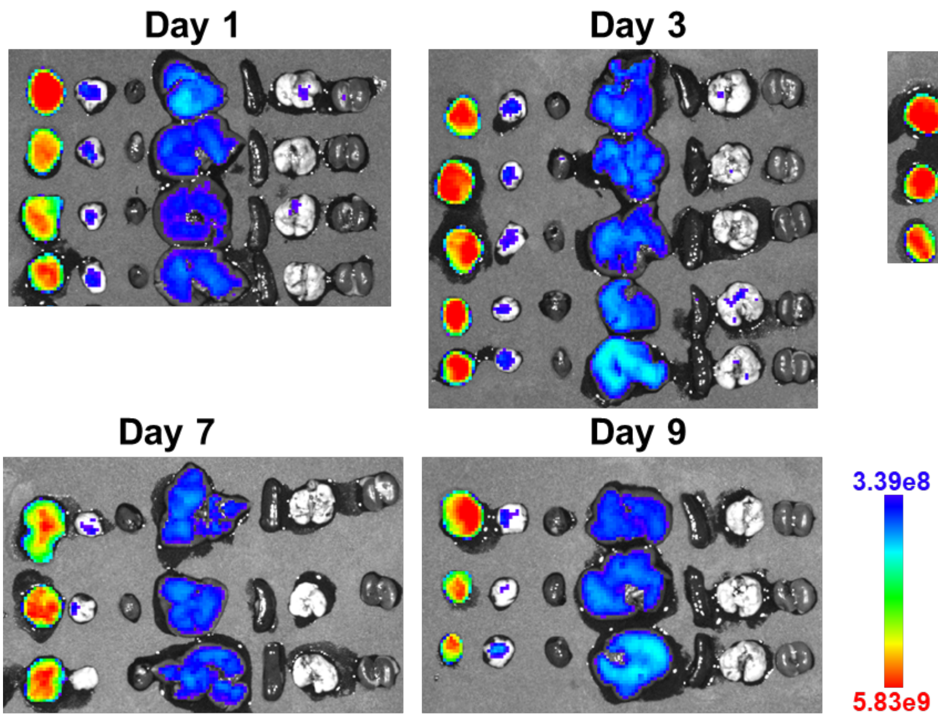


**Figure S12.** Ex vivo fluorescence imaging of major organs in C57BL/6 mice bearing Panc02 subcutaneous tumour after 1, 3, 5, 7, 9 days postinjection of OEGMA-11 NPs (organs from left to right: tumour, pancreas, heart, liver, spleen, lung, kidney; Dose of NPs: 30 mg/kg).

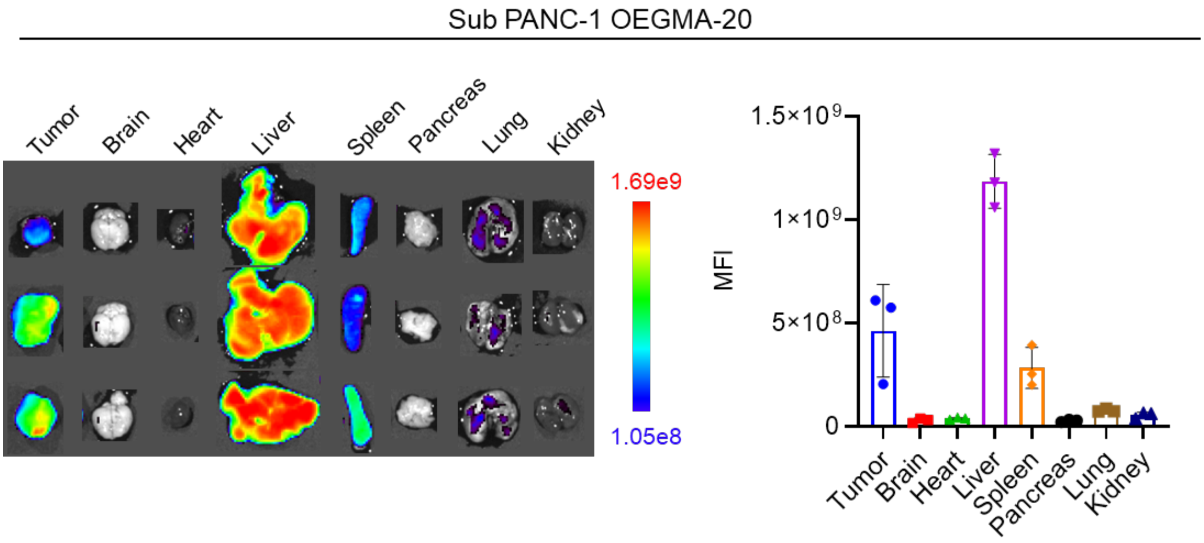


**Figure S13.** Ex vivo fluorescence imaging of major organs in BALB/c nude mice bearing PANC-1 subcutaneous tumour after 48 h postinjection of OEGMA-20 NPs (Dose of NPs: 30 mg/kg).

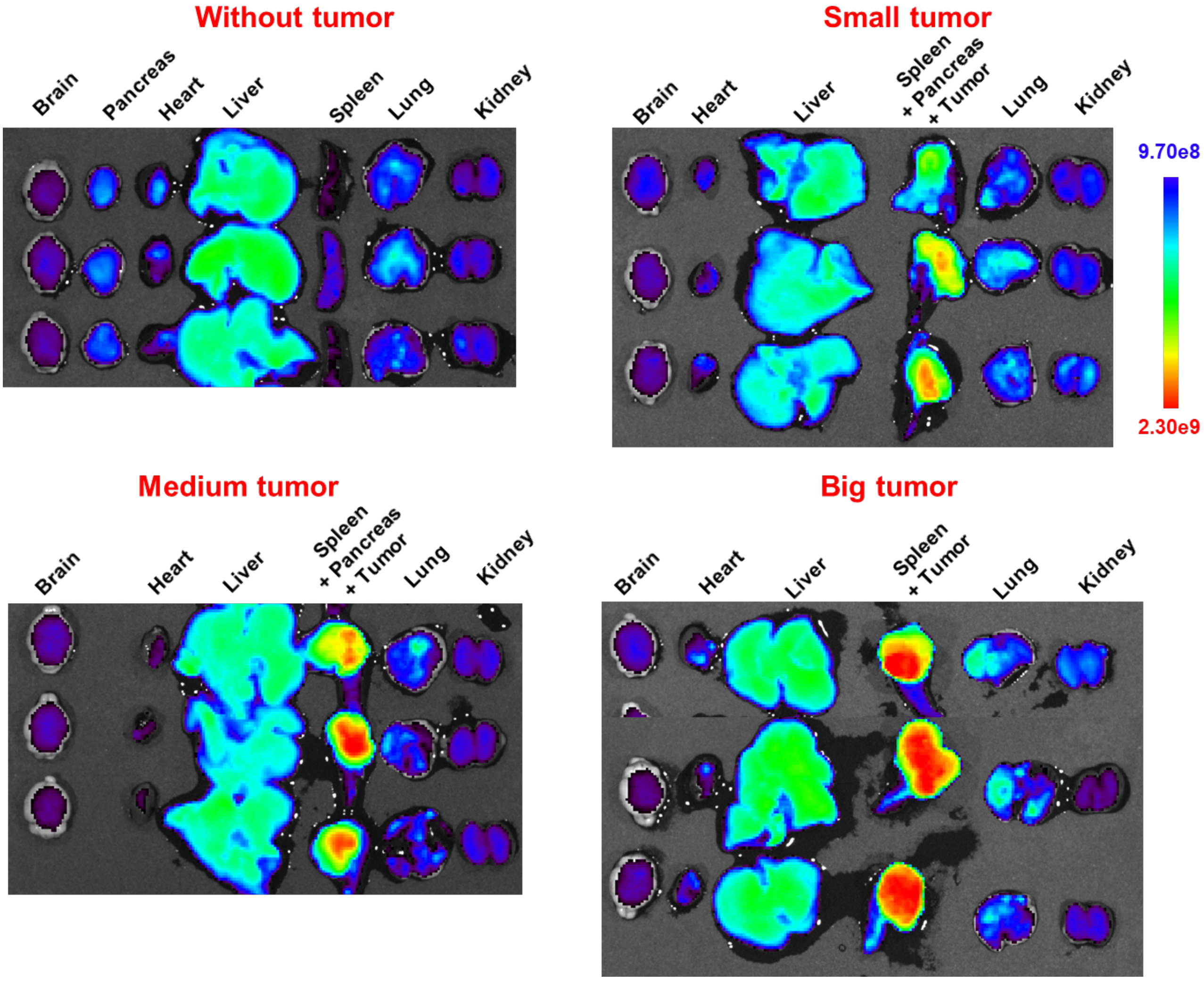


**Figure S14.** Ex vivo fluorescence imaging of major organs in BALB/c nude mice bearing PANC-1 orthotopic tumour (at different stages)/healthy mice after 48 h postinjection of OEGMA-11 NPs (Dose of NPs: 30 mg/g).

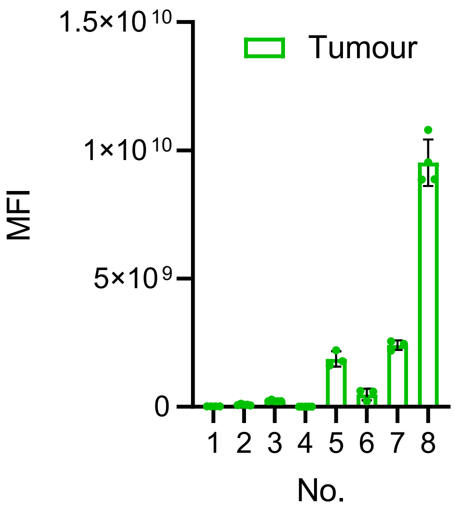


**Figure S15.** MFI of the tumour form Fig. **2b**, **3b**, **3j**, **4d** and **S12** (The number order is from **Fig. 5a**; n = 3 or 4 or 5).

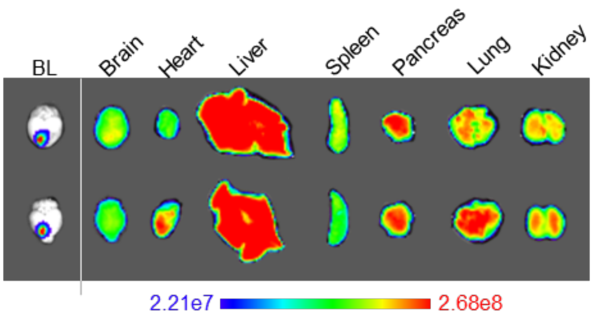


**Figure S16.** Ex vivo fluorescence imaging of main organs and U251- luc glioma at 48 h after injection of OEGMA-11 NPs (Dose of NPs: 30 mg/kg).

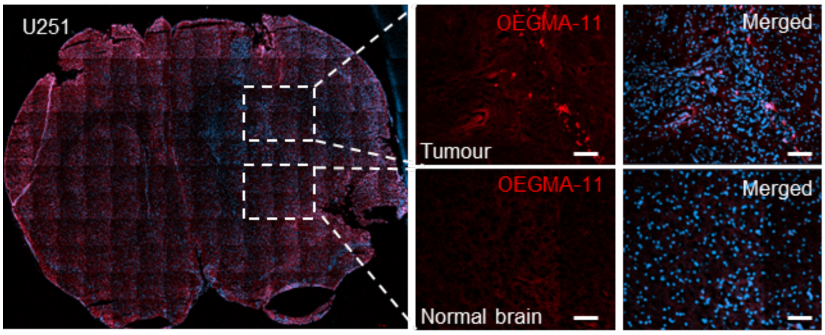


**Figure S17.** Fluorescent images of cryo-slice of brain of mouse bearing orthotopic U251-luc tumour after 48 h postinjection of OEGMA-11 NPs (Dose of NPs: 30 mg/kg, scale bar: 100 μm).

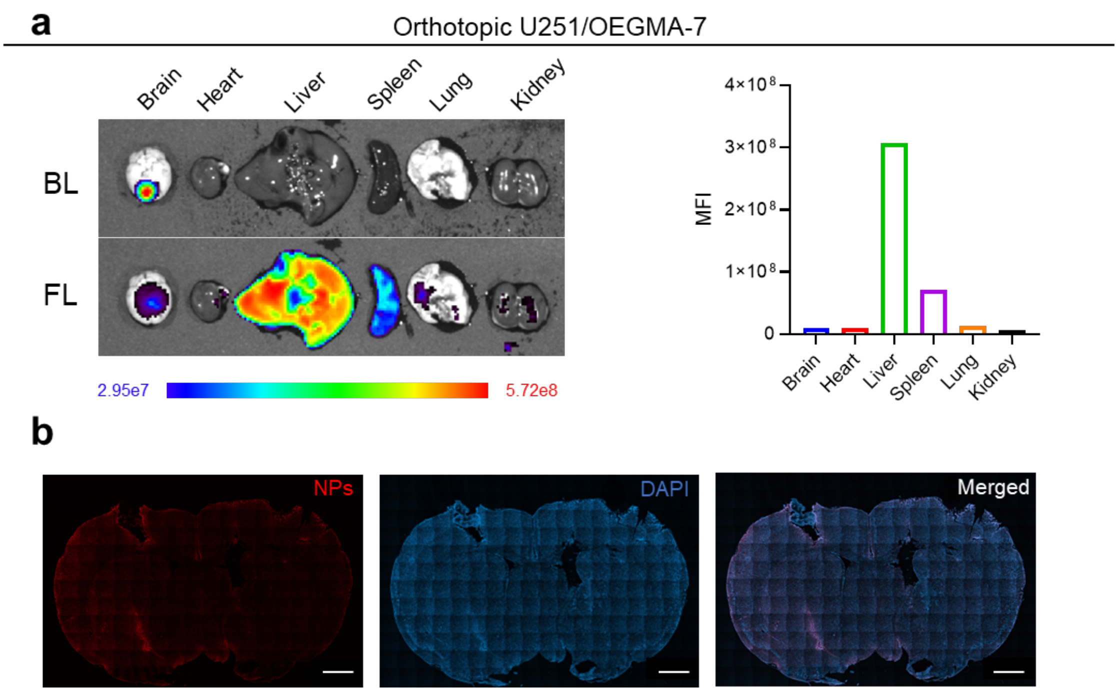


**Figure S18.** The biodistribution and penetration of OEGMA-7 NPs. **a** Ex vivo fluorescence imaging of major organs in BALB/c mouse bearing U251-luc orthotopic tumour at 48 h administration of OEGMA-7 NPs (Dose of NPs: 30 mg/kg). **b** Fluorescence images of cryo-slice of brain of mouse bearing orthotopic U251-luc tumour after 48 h postinjection of OEGMA-7 NPs (scale bar: 1000 μm).

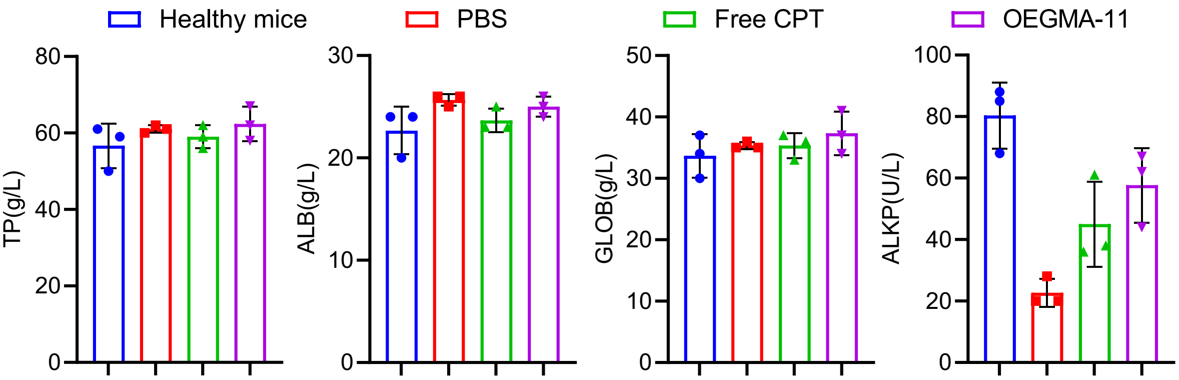


| Groups | Healthy mice | PBS | Free CPT | OEGMA -11 |
|---|---|---|---|---|
| CREA (μmol/L) | < 9 | < 9 | < 9 | < 9 |

**Figure S19.** Serum chemistry analysis of healthy mice and PANC-1 tumour bearing mice treated with PBS, free CPT and OEGMA-11 NPs (CPT-equivalent dose of 10 mg/kg, five doses every five days. Abbreviations: TP, total protein; ALB, albumin; GLOB, globulin; ALKP, alkaline phosphatase; CREA, creatinine).

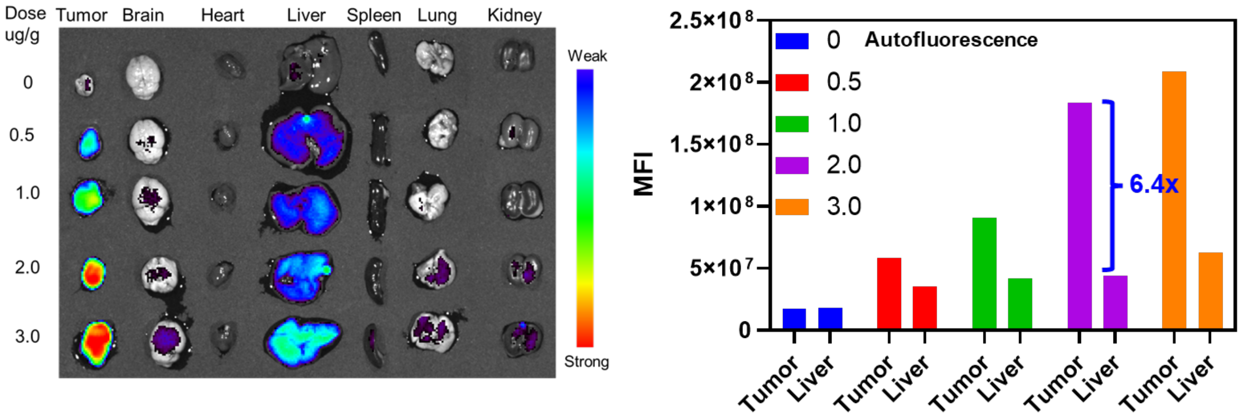


**Figure S20.** Ex vivo fluorescence imaging and MFI of major organs after 48 h postinjection of OEGMA-11 NPs with different doses in Panc02 tumour model.

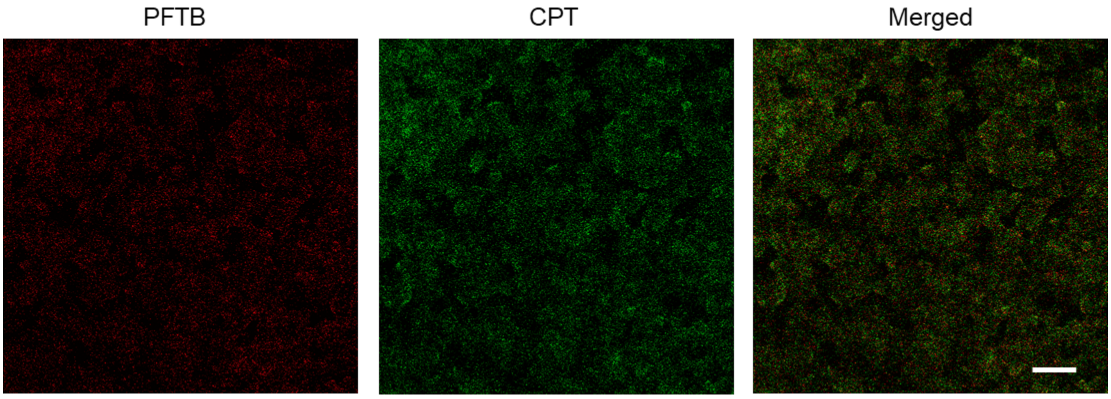


**Figure S21.** Fluorescence images of frozen slice of Panc02 tumour from mouse after 48 h postinjection of OEGMA-11 NPs (scar bar: 100 μm; Red and green fluorescence from backbone PTFB and CPT of OEGMA-11 NPs.).

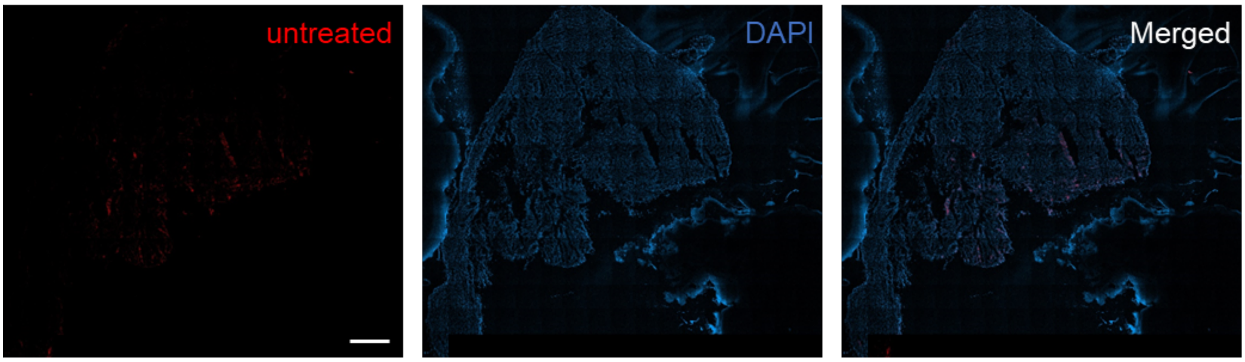


**Figure S22.** Fluorescence images of cryo-slice of 4T1 tumour injected with PBS at 48 h (scar bar: 1000 μm).

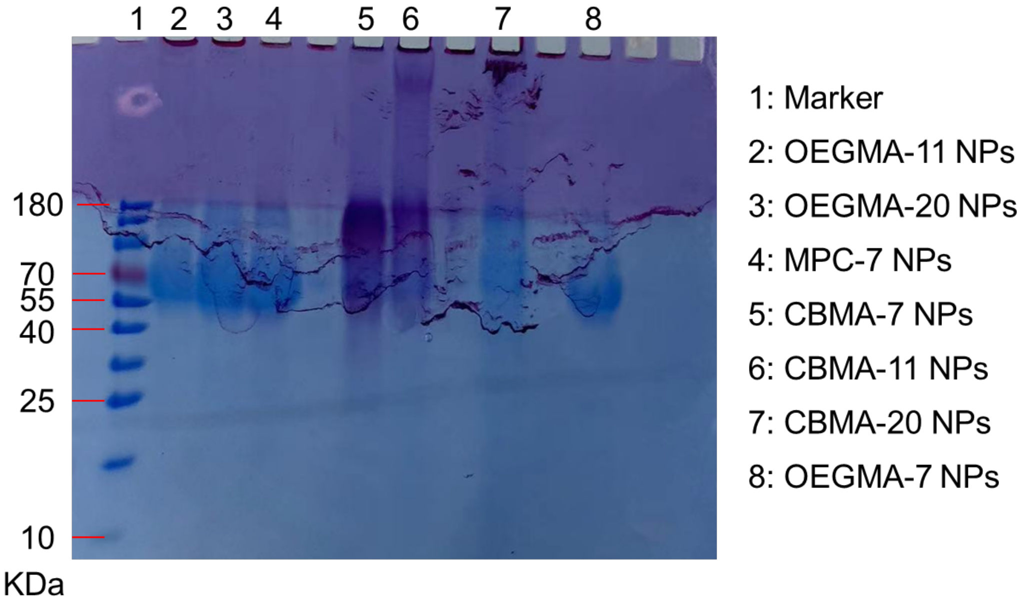


**Figure S23.** Sodium dodecyl sulphate-polyacrylamide gel electrophoresis (SDS-PAGE) of protein corona on the surface of NPs after incubation with mourse serum.

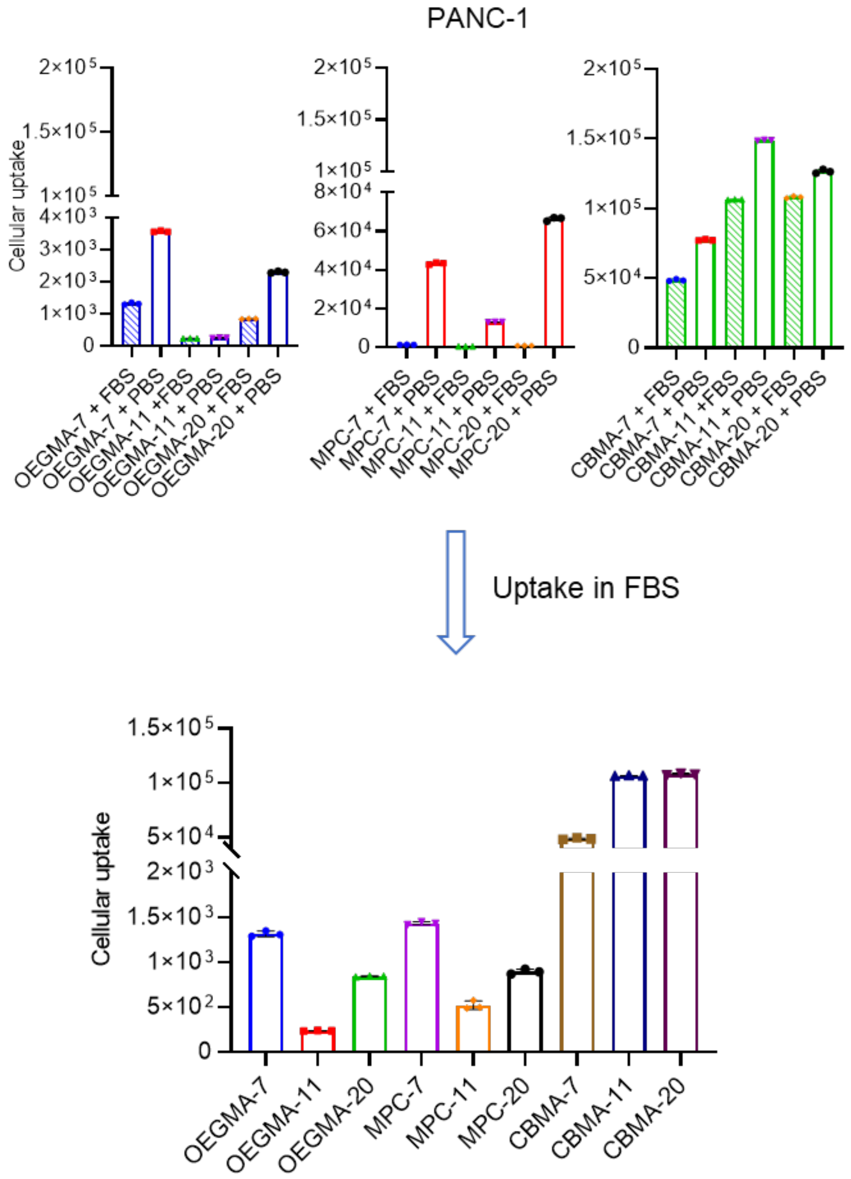


**Figure S24.** Flow cytometric analysis of the cellular uptake of PANC-1 cells treated with NPs in FBS/FBS-free medium for 4 h.

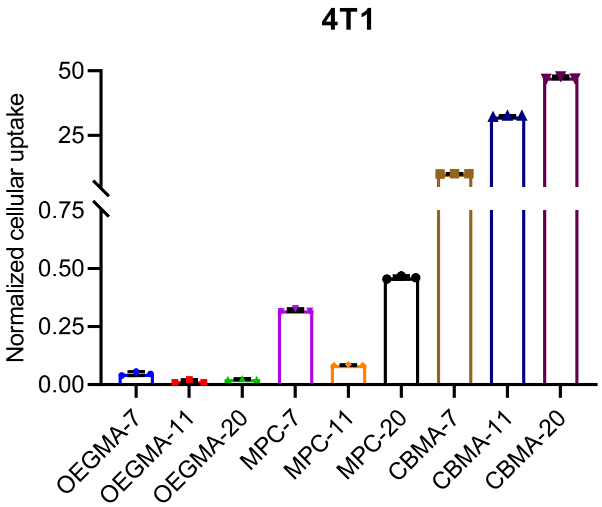


**Figure S25.** Flow cytometric analysis of the cellular uptake of 4T1 cells for 4 h.

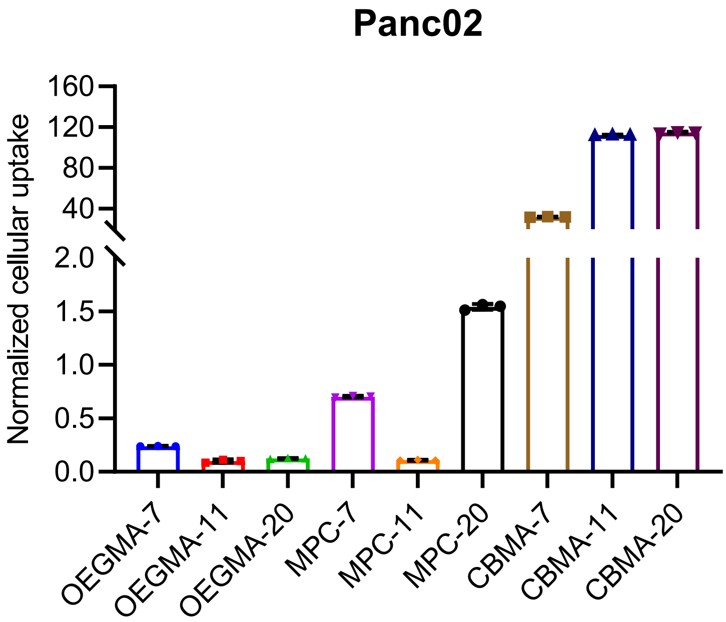


**Figure S26.** Flow cytometric analysis of the cellular uptake of Panc02 cells for 4 h.

**Table S1.** In vivo examples of tumour-targeting efficiency of nanomedicines in mouse tumour models.

| No. | nanomedicine platform | Active ingredients | Size (nm) | Surface potential (mV) | Blood circulation time (h) | Tumour model | Tumour-targeting efficiency |
|---|---|---|---|---|---|---|---|
| **1**[5] | Polymer | Doxorubicin/ ulixertinib | 9.0 | 1.0~2.8 | < 0.5 | Sub HT-29 | T/L < 1.0 |
| **2**[6] | Polymer | Resiquimod | 10.9 | / | 6.8 | Sub MC38 | T/L = 1.0 |
| **3**[7] | Polymer | OTX-015/AZD5153 | 15.2 | / | 1.8 | Sub 4T1 | T/L = 3.0 and 1.0 |
| **4**[8] | Polymer | Resiquimod | 50.0/70.0 | -13.0/-29.0 | / | Sub 4T1 | T/L = 2.0 |
| **5**[9] | Polymer | Camptothecin | 60.0 | -9.0 | < 0.5 | Sub HepG2 | T/L < 1.0 |
| **6**[10] | Polymer | 7-ethyl-10-hydroxyl camptothecin | 70.0 | -2.1 | 21.3 | Sub HepG2 | T/L = 8.0 |
| **7**[11] | Antibody+MSN | Camptothecin | 90.0 | -5.3 | / | Sub SK-BR3 | T/L = 9.0 |
| **8**[12] | CM+$SiO_2$ | / | 40.0 | -20.0 | 1.0 | Sub MCF-7 | T/L = 14.0 |
| **9** | Our polymer—OEGMA-11 | Camptothecin | 16.1 | - 7.0 | 27.0 | Sub 4T1 | T/L = 14.4 |
| | | | | | | Sub Panc02 | T/L = 11.8 |
| | | | | | | Sub PANC-1 | T/L = 9.1 |
| | | | | | | Ort PANC-1 | T/L = 4.7 |

CM: cell membrane; MSN: mesoporous silica nanoparticle; Sub: subcutaneous; Ort: orthotopic; T/L: MFI or content ratio of nanocarrier in tumour and liver.

**Table S2.** The molecular weight and polydispersity index ($M_w/M_n$) of **polymer 4** measured by GPC.

| **Polymer 4** | $M_n$ | $M_w/M_n$ | Repeating units (n) |
|---|---|---|---|
| n = 7 | 7600 | 1.4 | 7 |
| n = 11 | 11400 | 1.6 | 11 |
| n = 20 | 20400 | 1.2 | 20 |

**Table S3.** The half-lives of NPs in blood (unit: h).

| | **OEGMA** | **MPC** | **CBMA** |
|---|---|---|---|
| n = 7 | 1.9 | 4.9 | 0.9 |
| n = 11 | 27.0 | 0.8 | 0.5 |
| n = 20 | 0.9 | 0.6 | 0.45 |
